# Environmental phylogenetics supports a steady diversification of crown eukaryotes starting from the mid Proterozoic

**DOI:** 10.64898/2025.12.12.693929

**Authors:** Miguel M. Sandin, Phoebe Cohen, Hélène Morlon, Fabien Burki

## Abstract

Molecular clock and preservation of early microfossil assemblages suggest that eukaryotes were present and already diverse more than ∼1600 million years ago (Ma). Yet, the earliest identifiable eukaryotic crown group fossil only appeared around 1050 Ma, leaving a ∼600 million years gap in the Mesoproterozoic during which the evolution and diversification of early eukaryotes remain poorly understood. Here, we inferred a timeline of eukaryote evolution using molecular clock and birth-death diversification models, including the great diversity of environmental sequences to take into account the uncultured majority of microbial life. Our analyses, based on a unique dataset of 75,975 non-redundant Operational Taxonomic Units and 77 well-supported fossil calibrations, indicate a steady diversification of crown group eukaryotes during the Proterozoic after the Last Eukaryotic Common Ancestor (LECA). We show that Archaeplastida was one of the earliest diversifying supergroups and the most diverse throughout the Proterozoic, suggesting that the first successful plastid endosymbiosis gave Archaeplastida a measurable evolutionary advantage. Discoba, Amoebozoa and Rhizaria followed Archaeplastida in phylogenetic diversity throughout the Proterozoic, suggesting that crown eukaryotes were already thriving in this Era. These results contrast with the common view that the geologically stable Proterozoic Era experienced little eukaryotic evolutionary innovation, and indicate that it was already at the dawn of eukaryote evolution that all current supergroups were established.

## 2. Introduction

The origin of eukaryotes is a major evolutionary transition in the history of life that has shaped our planet in countless ways. Today, eukaryotes are found in virtually every environment on earth, ranging from tiny single-cells to the largest organisms to have ever lived. Although macro-organisms are often the most charismatic eukaryotes, it is the microbes that have dominated eukaryote evolution for most of their history (<u>Hedges et al. 2004</u>; <u>Agić and Cohen 2021</u>). These microbial eukaryotes continue to make up the vast majority of diversity across the eukaryotic tree of life, having evolved into an incredible diversity of forms, lifestyles, and functions (<u>Burki et al. 2020</u>). In recent years, we have learned a great deal about the origin of eukaryotes from the merging of Archaea and Bacteria (e.g., <u>Spang et al. 2015</u>; <u>Zaremba-Niedzwiedzka et al. 2017</u>; <u>Eme et al.</u> <u>2023</u>; <u>Donoghue et al. 2023</u>; <u>Kay et al. 2025</u>), but the early history of eukaryotes after their last common ancestor (LECA) remains poorly understood. In particular, we know little about the diversification dynamics of the main eukaryotic supergroups following their origin.

The oldest unambiguous eukaryotic fossils appeared ∼1750 million years ago (Ma) with acritarchs inferred to possess complex features such as a cytoskeleton and endomembrane system (<u>Lamb et al. 2009</u>; <u>Agić et al.</u> <u>2017</u>; <u>Javaux, 2025</u>). By ∼1640 Ma, eukaryotes were taxonomically more diverse corresponding to simple multicellular and possibly photosynthetic forms in shallow marine environments (<u>Riedman et al. 2023</u>; <u>Liu et al.</u> <u>2023</u>; <u>Miao et al. 2024</u>). While the acritarch fossil record has become richer and informative about the deep history of eukaryotes, it can only serve as a minimum time bound for the origin of total eukaryotic clade (i.e., stem and crown groups). Beyond the morphological fossil record, size-based ecosystem modeling suggests a complex eukaryotic community from ∼1700 Ma onwards, composed of heterotrophs, autotrophs, and even mixotrophs (<u>Eckford-Soper et al. 2022</u>). In addition, the rock record points to abundant sterol precursors of eukaryotic origin (protosteroids and ursteroids) during the late Paleoproterozoic and Mesoproterozoic (<u>Brocks</u> <u>et al. 2023</u>). However, it was not until the late Mesoproterozoic and early Neoproterozoic that undisputed fossil representatives of extant eukaryotic groups (crown groups) appeared. These fossils famously include Bangiomorpha, commonly accepted as a 1050 Ma red alga (<u>Gibson et al. 2018</u>), and the 950 Ma green algal *Proterocladus* (<u>Tang et al. 2020</u>), both members of the photosynthetic supergroup Archaeplastida. Therefore, there is a ∼600 million year gap in the fossil record between the origin of the total group eukaryotes and the ecological rise of the first crown groups, during which time the diversification dynamics of eukaryotes remains unknown.

Complementary to paleontology, molecular clock analyses and phylogenetic diversification models allow us to infer the divergence times and diversification of species across phylogenetic clades, particularly in groups with poor fossil record. Current estimates based on phylogenomics and advanced molecular clock analyses indicate that crown eukaryotes diversified ∼1850-1600 Ma (<u>Parfrey et al. 2011</u>; <u>Betts et al. 2018</u>; <u>Kay et al. 2025</u>), or possibly earlier (<u>Strassert et al. 2021</u>), in good agreement with the Paleoproterozoic fossil record reporting complex eukaryotic (stem or crown) communities. While rich in genomic data, these molecular clock analyses suffer from a relatively poor representation of the massive lineage diversity of microeukaryotes that has been revealed by environmental sequencing (<u>Burki et al. 2021</u>). This environmental diversity has not yet been used to study diversification patterns across all eukaryotes. Indeed, the few phylogenetic diversification analyses that have used it focused mostly on specific taxonomic groups such as diatoms (<u>Lewitus et al. 2018</u>) or ciliates (<u>Da</u> <u>Silva Costa et al. 2021</u>). As a consequence, we know little about the diversification history of early eukaryotes. While estimating diversity dynamics from molecular phylogenies remains a subject of debate (<u>Nee et al. 1994</u>; <u>Paradis 2004</u>; <u>Rabosky 2010</u>; <u>Beaulieu and O’Meara 2015</u>; <u>Rabosky 2016</u>; <u>Mitchell et al. 2018</u>), diversification models have greatly improved (<u>Rabosky et al. 2017</u>; <u>Helmstetter et al. 2022</u>; <u>Morlon et al. 2024</u>), for example showing success at estimating bacterial diversification over geological time for which the fossil record is sparse and cryptic (<u>Louca et al. 2018</u>).

Here, we used a molecular approach to studying the diversification dynamics of all major eukaryotic groups from the Proterozoic until today. We built a unique phylogenetic dataset of 75,975 non-redundant Operational Taxonomic Units (OTUs), integrating 77 well-supported fossil calibrations and information from the large environmental diversity with long-read metabarcoding data of the near-full eukaryotic ribosomal operon (18S-28S rDNA) and curated 18S rDNA sequences. Our analyses provide phylogenetic evidence of the complexity of eukaryotic communities during the Mesoproterozoic and show that crown group eukaryotes were taxonomically diverse nearly 800 million years before direct evidence of morphological diversification in the fossil record. These results suggest that crown group eukaryotes were not only present but diversifying and therefore active players in eukaryotic communities ever since their last common ancestor. Altogether, this study brings a novel and important molecular perspective in eukaryotic paleoecology and complements interpretation of the fossil record.

## 3. Results

### 3.1. Timetree of eukaryotes including the vast environmental diversity

In order to include as much diversity as possible in our analyses, we took advantage of the very large collection of ribosomal DNA (rDNA) available for microbial eukaryotes (protists), focusing on environmental sequences. Our dataset combines long-read metabarcoding data of the 18S-28S rDNA, which provide improved phylogenetic signal over classical short-read data, with available 18S rDNA reference sequences to obtain 75,975 curated non-redundant OTUs spanning the broad eukaryotic diversity. Topological constraints taken from a consensus of published phylogenomic studies were applied to fix ambiguous deep nodes using rDNA alone (**<u>Fig. S1</u>**). A range of 32 phylogenetic trees were reconstructed using alternative alignments and evolutionary models to take into account phylogenetic uncertainties (**<u>Fig. S2</u>**).

Given existing doubts on the position of the root of the eukaryotic tree, we tested rooting on Discoba or Amorphea, corresponding to two main alternative roots for the eukaryotic tree (<u>Cerón-Romero et al. 2022</u>; <u>Al</u> <u>Jewari and Baldauf 2023</u>; <u>Williamson et al. 2025</u>). The trees were time-calibrated using a main set of 77 diverse accepted fossil calibrations (**<u>Table S1</u>**; **<u>Fig. S3</u>**). The estimated dates, summarized as Lineages Through Time (LTT) plots, were largely consistent across the 32 timetrees (see **Fig. 1A** for the best log likelihood scoring tree). The root position had a minimal effect on the overall ages throughout the trees (**<u>Fig. S4</u>**), with a median root age of 1776 Ma (and maximum and minimum medians of 1897 and 1670 Ma, respectively) for the Discoba root (**Fig. 1B**) and 1773 Ma (1934-1703 Ma) for the Amorphea root (**Fig. 1C**). Below we arbitrarily focus on the Discoba-rooted trees for all subsequent results. Discoba was the first supergroup to diversify at 1594 (1726-1471) Ma, followed by Amoebozoa (1354; 1493-1257 Ma) and shortly after Archaeplastida (1343; 1421-1266 Ma) (**Fig. 1B**, yet see **Fig. 1C** and **<u>Fig. S4</u>** for different rooting scenarios and Highest Posterior Densities). All the other supergroups originated within a ∼200 million years span during the Mesoproterozoic starting ∼1275 Ma.

**Figure 1.**
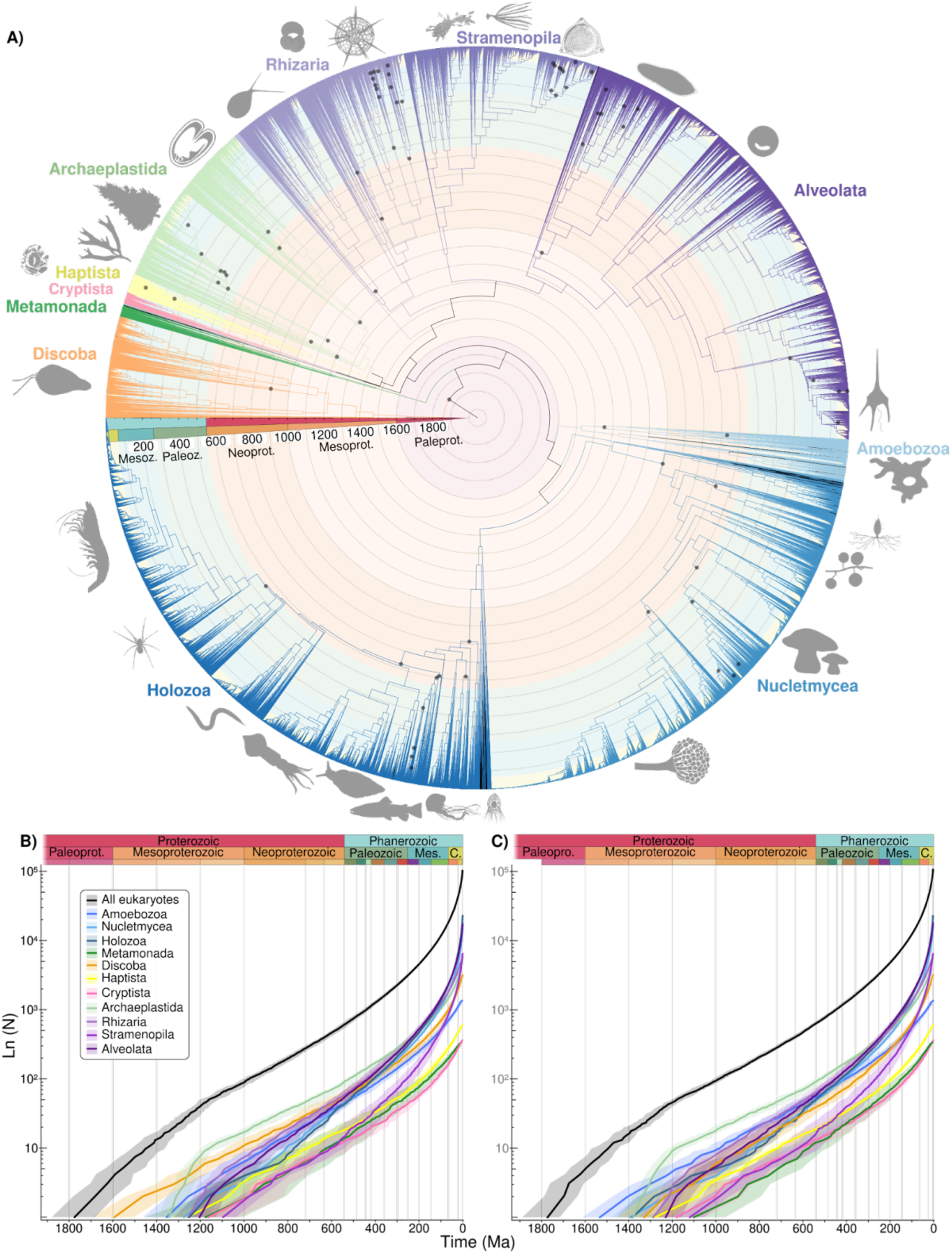
**A) A time-calibrated phylogenetic tree of extant eukaryotes**, rooted in Discoba, containing 71138 non-redundant 99% OTUs of the rDNA (5794 aligned positions) selected from the total 32 time-trees based on the highest likelihood. Coloured branches represent different eukaryotic supergroups (only those with more than 300 tips are named) and background colour different geological eras. Black dots on nodes highlight calibrated nodes. Silhouettes represent major branches of morphologically described diversity for visual guidance (e.g.; The fish on the lower left, *Salmo trutta*, represents all Chordate lineage, from other fishes and sharks to humans, birds and amphibians). **B and C) Summary of the Lineages Through Time (LTT) plot of all 32 time-calibrated trees**, rooted at Discoba (**B**) and Amorphea (**C**). Lines represent the median among all LTT and shaded areas represent the 90% centered percentile.

The impact of disputed fossil calibrations on our inferred dates was assessed by using two other calibration sets (see ‘Material and Methods’ for a rationale on the choice of the fossil calibrations). The effect of these uncertain fossils was variable depending on their age and taxonomic assignments. The Proterozoic *Rafatazmia* fossil, initially attributed to crown Rhodophyta (<u>Bengtson et al. 2017</u>) but later questioned (<u>Carlisle et al. 2021</u>; <u>Gibson et al. 2018</u>), had a large effect by pushing back the first diversification of eukaryotes at 2054 (2104-1967) Ma (**<u>Fig. S5</u>**). Not considering the calibration corresponding to the first diversification of diatoms, as recently argued (<u>Bryłka et al. 2023</u>), had no significant changes (**<u>Fig. S6</u>**).

### 3.2. Estimating total diversity of eukaryotes

Phylogenetic diversification analyses require an estimate of the sampled fraction of the hypothetical total diversity of the taxonomic groups under scrutiny to reliably estimate speciation and extinction rates (<u>Stadler</u> <u>2009</u>; <u>Helmstetter et al. 2022</u>). We sampled the largest available eukaryotic diversity while maintaining enough phylogenetic signal, from high-rank to species-level taxa including both morphologically described species and the uncultured majority of environmental sequences. We implemented two different approaches relying on species abundance distributions to estimate the fraction of the total eukaryotic diversity included in our data (see Material and Methods). On average, our phylogenetic trees contained between 77% and 50% of the hypothetical total eukaryotic diversity as defined by OTU clustering at 99% of the 18S rDNA (**<u>Fig. S7</u>**). However, this diversity fraction is not uniformly distributed across the tree. While groups such as Nucletmycea and Holozoa were relatively well sampled, others such as Rhizaria, Alveolata and Cryptista were less well represented. In general, these groups also have a much lower proportion of morphologically described species than environmental OTUs, as compared to groups such as Holozoa (**<u>Fig. S8</u>**).

### 3.3. Diversification of eukaryotes

While insights into the phylogeny and early evolution of eukaryotes have greatly improved in the last 20 years, the patterns of diversification across the eukaryotic tree, especially during the Proterozoic, have been far less explored. Using our taxon-rich dataset, we analyzed the diversification of each supergroup using the ClaDS phylogenetic birth-death model (<u>Maliet et al. 2019</u>). This model accounts for rate heterogeneity across lineages, making it suitable for large scale phylogenetic diversification analyses. The newer implementation of ClaDS that uses data augmentation (<u>Maliet and Morlon 2022</u>) provided us with estimates of Diversity Through Time (DTT, **Fig. 2**, see **<u>Fig. S9</u>** for other rooting and diversity fraction scenarios), as well as mean speciation (λ) and extinction rates (μ) through time, from which net diversification rates can be estimated (r=λ-μ, **<u>Fig. S11</u>**).

**Figure 2.**
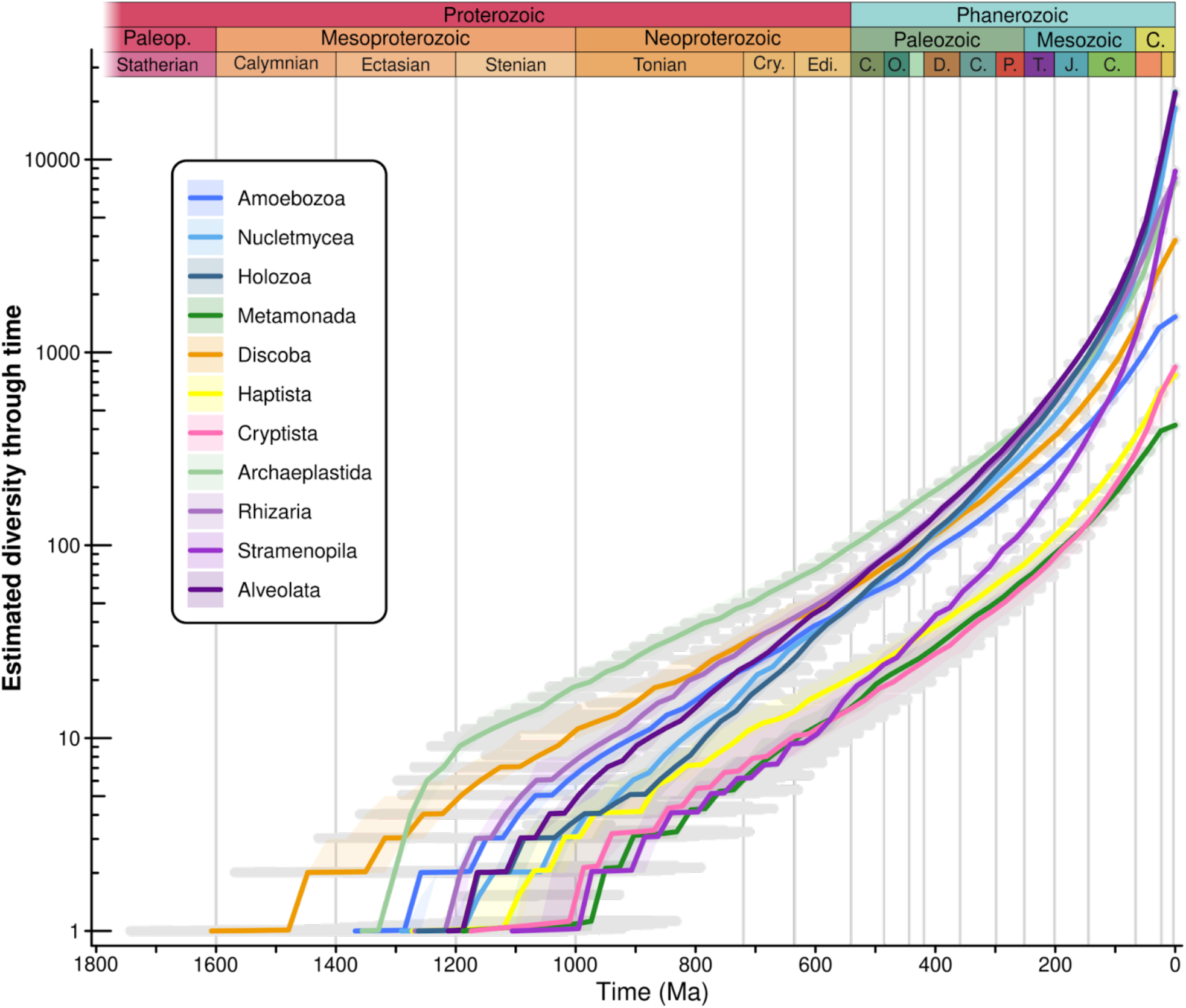
**Summary of the estimated Diversity Through Time (DTT) plots obtained with ClaDS**, for the Discoba rooted scenario assuming the higher range of diversity fractions (see **<u>Fig. S9</u>** for the Amorphea rooted scenario and the other diversity fractions). Lines represent the median and shaded areas the 90% Highest Posterior Density (HPD), respectively, of all independent DTT. Gray horizontal lines represent the 90% HPD of the time scale.

Archaeplastida and Rhizaria accumulated diversity rapidly shortly after their origin, before slowing down in the late Mesoproterozoic and Neoproterozoic and later increasing again (**Fig. 2**). Archaeplastida experienced the fastest early accumulation of lineages after their origin at 1343 (1391-1276) Ma and for the longest time period, resulting in the highest diversity between ∼1300 Ma until the early Mesozoic (**Fig. 2**). This rapid accumulation of lineages is also seen within the two most specious archaeplastid groups (Chloroplastida and Rhodophyta) (**<u>Fig. S10</u>**), suggesting a general trend in Archaeplastida. The supergroups other than Archaeplastida, and to some extent Rhizaria, first accumulated diversity relatively slowly but steadily starting in the Proterozoic, before an acceleration towards present (**Fig. 2**).

The trends in diversification rates are remarkably consistent across groups showing a steady increase towards the present (**<u>Fig. S11</u>**). This is reflected in the ClaDS trend parameter α, and heterogeneity parameter σ, which together predict that daughter lineages have on average a higher speciation rate than their parents (m>1, **<u>Fig.</u> <u>S12</u>**). Although extinction rate estimates are overall low compared to speciation, most differences in diversification rates are due to differences in extinction rates, with supergroups with a lower OTU richness (i.e., Cryptista, Haptista and Metamonada) showing some of the highest extinction rates (**<u>Fig. S11</u>**).

### 3.4. Speciation rate shifts across the tree

A shift increase of the speciation rate on specific branches of the tree can for example result from key innovations, specific adaptations, or variations in biotic and/or abiotic conditions that promote speciation. Similarly, a shift decrease can arise for example if changes in environmental conditions impede speciation. We estimated speciation rate shifts using BAMM (see Material and Methods) for all different supergroups within all 32 timetrees independently. At face value, most identified shifts in the speciation rate occurred closer to the present (**<u>Fig. S13</u>**), as expected based on the fact that there are more lineages, cumulating more evolutionary time, close to present in phylogenetic trees. However, when correcting the number of shifts by the cumulative time during which they had the opportunity to occur, measured as the total branch-length within geological time periods, the frequency of speciation rate shifts is instead slightly reduced towards present in all supergroups (**Fig. 3**, see **<u>Fig. S14</u>** for other rooting and diversity fraction scenarios). The most marked decrease in the frequency of shifts occurs from the Mesoproterozoic to the Neoproterozoic, going from a median 4.9·10^-4^ shifts per-lineage per-Myr to 1.74·10^-4^. After that, the frequency of shifts stays relatively constant from the Neoproterozoic through the Paleozoic. In most supergroups we inferred a higher proportion of speciation rate increases than decreases (**Fig. 3**, **<u>Fig. S14</u>**), further supporting the tendency for an increase in diversification rates through time found with ClaDS, and suggesting that diversity “begets” rather than “impedes” more diversity. In contrast, Rhizaria stands out as experiencing the highest proportion of diversification rate decays during the Proterozoic and Phanerozoic, followed by Stramenopila and Alveolata (**Fig. 3**, **<u>Fig. S14</u>**). Overall, the mild decline in rate shifts over time suggests that lineages experienced relatively more key innovations or adaptations that modified their speciation rates early in their evolutionary history.

**Figure 3.**
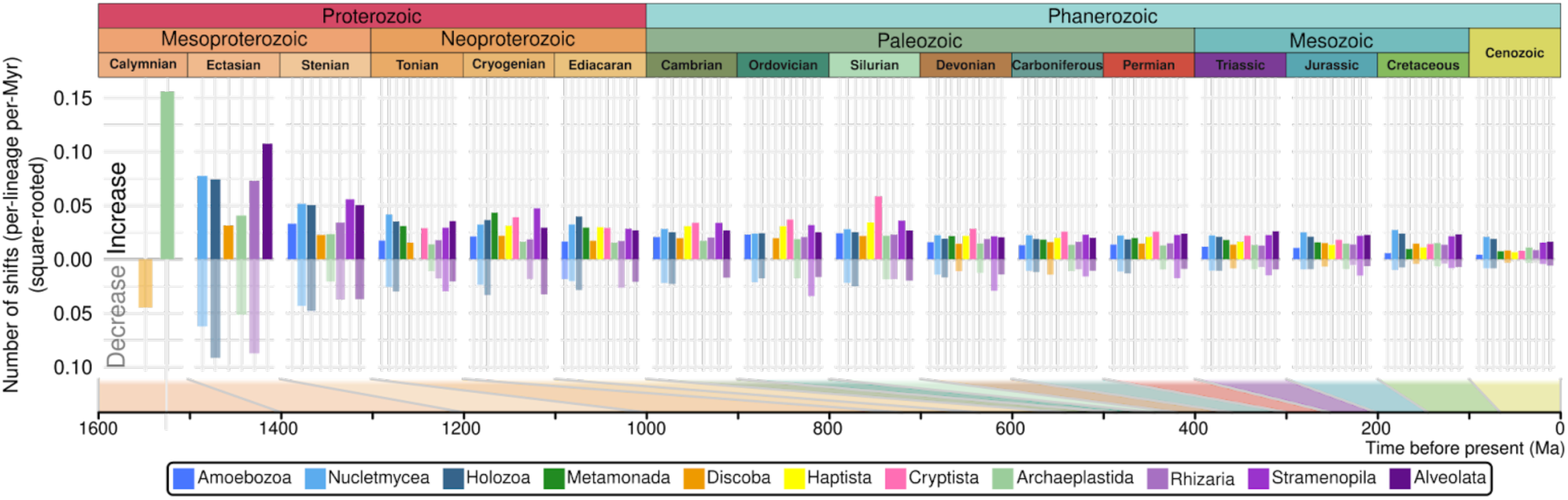
**Proportion of shifts in speciation rate across geological time**, under the Discoba rooted scenario and assuming the higher range of diversity fractions (see **<u>Fig. S14</u>** for the Amorphea rooted scenario and the lowest diversity fractions). ‘Increase’ indicates a shift towards a higher speciation rate and ‘decrease’ indicates a shift towards a lower speciation rate. The proportion of shifts was calculated by dividing the total number of shifts per time period (**<u>Fig. S13</u>**) by the total branch-length within the given time period.

## 4. Discussion

Here we have integrated the phylogenetic information from both environmental and reference rDNA sequences into the most taxon-rich molecular clock and diversification analysis across eukaryotes to date. Our analyses place the Last Eukaryotic Common Ancestor (LECA) at ∼1775 Ma, in agreement with the first unequivocal eukaryotic fossils (<u>Lamb et al. 2009</u>; <u>Agić et al. 2017</u>; <u>Javaux 2025</u>) and other molecular dating analyses of eukaryotic origin (<u>Parfrey et al. 2011</u>; <u>Eme et al. 2014</u>; <u>Betts et al. 2018</u>; <u>Mahendrarajah et al. 2023</u>; <u>Kay et al.</u> <u>2025</u>; **<u>Fig. S15</u>**). Including older but debated calibrations for crown eukaryotes alters the timing of LECA. Specifically, if Rhodophytes are as old as 1600 Ma as implied by the multicellular fossil *Rafatazmia* (<u>Bengtson et</u> <u>al. 2017</u>), our analyses shifted the origin of LECA to the mid Paleoproterozoic (∼2054 Ma). Recent phylogenomic analyses indicated a Paleoproterozoic appearance of Archaeplastida (<u>Strassert et al. 2021</u>) even without the *Rafatazmia* calibration (<u>Sánchez-Baracaldo et al. 2017</u>; <u>Yang et al. 2023</u>; <u>Bowles et al. 2024</u>), and thus an earlier origin of LECA and some crown eukaryotes is not without molecular support.

Beyond *Rafatazmia*, whose age and taxonomic assignment to a crown group have been questioned (<u>Gibson et</u> <u>al. 2018</u>; <u>Carlisle et al. 2021</u>), rich and complex assemblages of eukaryotes have been described from the late Paleoproterozoic, including both single-cell fossils from at least ∼1750 Ma (<u>Javaux 2025</u>) and simple multicellular organisms at ∼1640 Ma (<u>Riedman et al. 2023</u>; <u>Liu et al. 2023</u>; <u>Miao et al. 2024</u>). These fossil data suggest that eukaryotes were present and well established by the late Paleoproterozoic, but a major unanswered question in paleobiology is whether these early fossils may be unidentified members of crown eukaryote groups, thus bridging the gap between LECA (inferred here at 1775 Ma) and the ecological rise of extant supergroups during the late Proterozoic. A temporal offset between the fossil record and molecular dating is to be expected given the sparse Proterozoic fossil and rock record (<u>Peters et al. 2022</u>), leading to an amplified “Sppil-Rongis” effect (<u>Signor and Lipps 1982</u>). This effect states that incompleteness in the fossil and rock record will necessarily mean that the oldest known fossil in a clade will always be younger than the true clade origination date. This effect is well characterised for much more recent animal and plant fossils (<u>Foster et</u> <u>al. 2017</u>; <u>Barba-Montoya et al. 2018</u>; <u>Álvarez-Carretero et al. 2022</u>), but is likely even more pronounced for older protist forms that may lack fossilized and/or recognizable structures. While our molecular timing and diversification analyses inferred from only extant taxa cannot place enigmatic acritarchs with a eukaryotic phylogeny, they can help us better understand the evolutionary dynamics of the major eukaryotic groups following their appearance.

During the Mesoproterozoic and early Neoproterozoic, clearly identifiable crown group fossils or crown group biomarkers (i.e., sterols) are lacking. Yet, our results arguably show that most eukaryotic supergroups originated during this time. As for the “Sppil-Rongis” effect, sterol record may also suffer from taphonomic processes (<u>Cohen and Kodner 2022</u>); it is therefore possible that crown group eukaryotes were confined to specific environments where they diversified, but left limited fossil and biomarker evidence behind at a global scale. Coastal environments, for example, were likely oxygenated enough to sustain the development of eukaryotes since the Great Oxidation Event ca. 2.4 Ga (<u>Lyons et al. 2014</u>), and consistently supported high richness, morphological complexity and even habitat specialization (<u>Javaux et al. 2001</u>; <u>Javaux and Knoll 2017</u>). These relatively stable and productive coastal ecosystems may have provided the necessary conditions for both the origin of eukaryotes and their initial gradual diversification into the distinct supergroups reported in our analyses. However, speciation rates of the different supergroups remained relatively low during the Proterozoic compared to more recent times. Such slow, but substantial, initial diversification might be related to competition with other contemporary lineages, including stem and crown species, as previously suggested (<u>Eckford-Soper et al. 2022</u>; <u>Liu et al. 2023</u>). Similar competition patterns between more recent kin lineages have been reported, where angiosperms ultimately outcompeted conifers from tropical dominance over the last ∼350 million years (<u>Condamine et al. 2020</u>).

One supergroup that differs from the general pattern of slow but steady initial diversification is Archaeplastida. Our analyses indicate that Archaeplastida experienced a fast and early diversification throughout the Proterozoic, dominating the phylogenetic diversity of crown eukaryotic communities during this period. We postulate that this early success of Archaeplastida is the result of the cyanobacterial endosymbiosis in an ancestor of this group, responsible for establishing the first photosynthetic organelles (plastids) and thus for giving eukaryotes the transformative capacity to use light to produce chemical energy (<u>Bhattacharya and</u> <u>Medlin 1995</u>; <u>Bhattacharya et al. 2004</u>). The rapid diversification of Archaeplastida is also seen in the fossil record, with many putative archaeplastid fossils identified during the late Paleoproterozoic (<u>Liu et al. 2023</u>; <u>Miao et al. 2024</u>), probably representatives of stem Archaeplastida (<u>Miao et al. 2024</u>). The early success of Archaeplastida reached far beyond the limit of this group, ultimately impacting the evolution of all photosynthetic eukaryotes. From archaeplastid algae, photosynthetic organelles were transferred by several rounds of eukaryote-to-eukaryote endosymbioses (<u>Strassert et al. 2021</u>) to other crown groups that we infer were slowly diversifying during the Proterozoic. Supergroups such as Cryptista, Haptista and to a lesser extent Alevolata and Stramenopila show relatively small pulses of diversification rates throughout the Proterozoic, perhaps a phylogenetic record of the acquisition of plastids in these groups.

Archaeplastida might have also impacted early on non-photosynthetic eukaryotic groups. Our analyses reveal high phylogenetic diversity in Rhizaria and Discoba (or Amoebozoa, depending on the rooting scenario) throughout the Proterozoic, which are supergroups that use a wide variety of heterotrophic strategies from active predation to parasitism. Recent fossil evidence suggests increasingly complex ecological dynamics, including ectosymbiosis by 1000 Ma (<u>Tang et al. 2021</u>) and eukaryovory likely even as far back as ∼1750 Ma (<u>Javaux 2025</u>). Ovoid perforations found in organic-walled assemblages from 1150-900 Ma were interpreted as traces of selective predation (Loron et al. 2018). Early biomineralized scales ca. 810 Ma were interpreted as defense against predation (<u>Cohen et al. 2017</u>), or direct evidence of predators themselves with the first testate amoebae at ca. 740 Ma (<u>Porter et al. 2003</u>; <u>Dumack et al. 2024</u>). It is therefore possible that the early diversification of the heterotrophic lineages Rhizaria, Discoba and Amoebozoa was spurred in part by the earlier expansion of Archaeplastida, perhaps used as prey along other members of the community, as well as the slow but inexorable oxygenation of the deep ocean (<u>Alcott et al. 2024</u>).

## 5. Conclusions

The early evolution of eukaryotes remains poorly understood, owing in part to the sparsity of the Proterozoic rock and fossil record (i.e. <u>Porter et al. 2025</u>), and difficulties in associating many early Proterozoic fossils with present day taxonomic groups. At the same time, molecular data has remained too scarce for the uncultured majority of microbial life to better represent the vast diversity of eukaryotes, at least for applying phylogenetic and diversification models. Here, we take a significant step toward resolving these challenges by integrating environmental diversity data with richer phylogenetic signals into a diversification framework spanning all major eukaryotic supergroups. Even so, our estimates based on this large dataset suggest that we may have captured as little as half of the total eukaryotic diversity, underscoring how much more hidden diversity awaits discovery. This is of course a limitation imposed by current environmental sampling, but this limitation is in part alleviated by the use of diversification methods that account for partial sampling. Our broadscale analyses indicate that crown group eukaryotes were not just present in the Proterozoic, but diversifying and probably actively engaging in ecological interactions that may have promoted further their diversification. Therefore, the geologically stable and fossil-poor billion-years-period spanning the late Palaeoproterozoic to early Neoproterozoic, sometimes referred to the “boring billion”, was in fact biologically exciting (<u>Mukherjee et al.</u> <u>2018</u>; <u>Javaux and Lepot 2018</u>; <u>Eckford-Soper et al. 2022</u>; <u>Mitchell and Evans 2024</u>) witnessing a period of steady diversification that ultimately resulted in the ecological rise of crown eukaryotes.

## 6. Material & Methods

All scripts, raw files, resulting phylogenetic trees and other resources needed for the replication of the analyses performed and presented in this study are publicly available at ZENODO (doi: 10.5281/zenodo.17901848) and explained in detail in <u>github.com/MiguelMSandin/EarlyDiverEuk</u>. In total, all successful analyses presented in this study were estimated to last ∼136.2 CPU years.

### 6.1. Dataset assembly

The Protists Reference Ribosomal database (PR2; <u>Guillou et al. 2013</u>; version 4.14) of the 18S ribosomal DNA (rDNA) was combined with a recently published dataset of the near-full length rDNA (18S + 28S rDNA; <u>Jamy et</u> <u>al. 2022</u>) obtained from long-read sequencing environmental samples through Circular Consensus Sequencing by Pacific Bioscience (referred as PacBio dataset from here on). In order to avoid redundancy, the 183 237 nuclear eukaryotic sequences of the PR2 database were clustered at 99% Operational Taxonomic Units (OTUs) with Mothur v1.44.1 (<u>Schloss et al. 2009</u>), resulting in 59 510 OTUs. The two datasets were compared for identical sequences with VSEARCH v2.14.1 (<u>Rognes et al. 2016</u>) and removed the shortest identical sequence to maximize phylogenetic information. In total, 59 154 sequences of the PR2 database and 16 821 from the PacBio dataset were assembled together into 75 975 non-redundant OTUs of the rDNA.

We have additionally considered the global dataset of the V4 hypervariable region of the 18S rDNA (EukBank: (<u>Berney et al. 2023</u>) to increase richness diversity into our analyses. In total, 335 662 metabarcodes of the V4 hypervariable region of the 18S rDNA gene were compared against our previously assembled dataset for similar sequences with VSEARCH v2.14.1 (<u>Rognes et al. 2016</u>). Of all metabarcodes, 10 717 (3.2%) were identically found in the previously assembled dataset, 40 286 (12.0%) were at least 99% similar, 99 565 (29.7%) were at least 97% similar and 95 350 (28.4%) were at most 80% similar. Given (i) the relatively high number of metabarcodes with low pairwise similarity, (ii) their low phylogenetic information (357.2 ± 60.2 average base pair length), (iii) the difficulties to analyze phylogenetic placement of distant metabarcodes at global eukaryotic scale (<u>Ewers et al. 2023</u>), (iv) the ambiguities on the nature of high-throughput short metabarcodes (i.e.; sequencing errors vs intragenomic variability: <u>Sandin et al. 2022</u>; <u>Greco et al. 2023</u>), and (v) a trade-off between sufficient diversity coverage and keeping computational efforts technically feasible, we decided not to include this dataset in our analyses.

### 6.2. Build initial constraint tree

Initial constraint tree (**<u>Fig. S1</u>**) was obtained from the EukProt database (<u>Richter et al. 2022</u>) and complemented with the latest phylogenomic studies within specific supergroups: Amoebozoa (<u>Kang et al. 2017</u>), Alveolata (<u>Tikhonenkov et al. 2020</u>), Archaeplastida and Cryptista (<u>Schön et al. 2021</u>; <u>Irisarri et al. 2022</u>; <u>Yazaki et al.</u> <u>2022</u>), Centrohelida and Haptista (<u>Burki et al. 2016</u>), Nucletmycea (<u>Li et al. 2021</u>), Opisthokonta (<u>Torruella et al.</u> <u>2015</u>), Rhizaria (<u>Irwin et al. 2019</u>), Stramenopila (<u>Derelle et al. 2016</u>), Telonemia (<u>Strassert et al. 2019</u>) and other orphans groups (<u>Brown et al. 2018</u>) such as Hemimastigophora (<u>Lax et al. 2018</u>) and Malawimonads (<u>Heiss et al. 2018</u>). When specific nodes showed contrasting topologies in different studies or low support, the given node was not resolved in the initial constraint tree and was left as a polytomy. Specific nodes relevant for fossil calibration were further resolved based on comprehensive phylogenomic and/or phylogenetic studies. These nodes are: Bilateria (<u>Telford et al. 2015</u>; <u>Dunn et al. 2014</u>) within Opisthokonta; Ciliophora (<u>Gao et al.</u> <u>2016</u>) and Dinoflagellata (<u>Janouškovec et al. 2017</u>) within Alveolata; Radiolaria (<u>Sandin et al. 2025</u>) and Foraminifera (<u>Pawlowski et al. 2013</u>) within Rhizaria; and Ochrophyta (<u>Di Franco et al. 2022</u>) within Stramenopila. The constraint tree was optimized over several phylogenetic analyses to avoid constraints generating polytomies or near-0 branches. Final constraint tree used for downstream analyses was obtained by replacing the given clade name (from **<u>Fig. S1</u>**) by all the sequences bearing such name in a common polytomyc node (described in detail in the github repository and available in ZENODO).

### 6.3. Progressive phylogenetic reconstruction of >75000 taxa

Phylogenetic analyses were performed in 3 steps over the 75975 non-redundant eukaryotic OTUs of the rDNA with the constraint tree built in the previous section:

-The first step was aimed to build a solid back-bone phylogenetic tree containing OTUs biologically relevant. Such OTUs were defined as representatives of at least 10 other sequences (for the PR2 database) or with at least 10 reads (for the PacBio database). Specific eukaryotic groups with a low representation in databases or environmentally scarce yet holding a valuable phylogenetic position were manually added to the previous selection of OTUs. These groups were Ancoracystida, Cephalochordata, Filasterea, Glaucocystophyceae, Gromia, Hemichordata, Katablepharidaceae, Malawimonadidae, Mantamonadida, Mesostigmatophyceae, Monothalamids, Noctilucophyceae, Palmophyllophyceae, Parabasalia, Placozoa, Pluriformea, Preaxostyla, Protalveolata, Rhodelphea, Synchromophyceae and Tubothalamea. The combined initial dataset contained a total of 11917 OTUs (11856 OTUs representative of at least 10 other sequences and 61 OTUs manually selected) and was automatically aligned using MAFFT v7.407 (<u>Katoh and Standley 2013</u>). The initial dataset was additionally reversed (with an in-house script ‘<u>fastaRevCom.py</u>’ available at the GitHub repository) and both forward and reverse files were aligned to account for possible misalignment issues. Aligned datasets were trimmed with trimAl v1.4.1 (<u>Capella-Gutiérrez et al. 2009</u>) using a 5% gap threshold (‘-gt 0.05’), resulting in 7123 and 7304 positions respectively for the forward and reverse alignments. In order to allow phylogenetic uncertainty, we applied 4 different phylogenetic analyses for both the forward and the reverse alignments using the tree generated in the previous step as a constraint topological tree of major nodes. Two thorough maximum likelihood searches were done in RAxML (<u>Stamatakis 2014</u>) under the GTR model of evolution (<u>Tavaré 1986</u>) and Gamma and CAT substitution rates optimization models over 100 bootstraps. The option ‘-D’ was used to stop maximum likelihood convergence criterion if the relative Robinson-Foulds distance between the trees obtained from two consecutive lazy SPR cycles is smaller or equal to 1%. Ten additional quick searches were obtained in RAxML-NG (<u>Kozlov et al. 2019</u>) with a GTR model of evolution and Gamma substitution rates optimization models. The 2 best scoring trees were kept for downstream analyses in order to allow a greater phylogenetic uncertainty of deep and ancient nodes. In total, 8 backbone phylogenetic trees were obtained at this step: RaxML GTR+Gamma, RaxML GTR+CAT and 2 runs in RaxML-NG for both the forward and reverse alignment. Resulting backbone phylogenetic trees of this step were processed to remove long branches using the function ‘prune’ from the package Bio.Phylo (<u>Talevich et al. 2012</u>); implemented in the in-house script ‘<u>treePruneOutliers.py</u>’). Briefly, long branches were determined by identifying outliers from a normal distribution (applying the generalized Extreme Studentized Deviate, gESD, method from; <u>Rosner 1983</u>).

-The second and third steps were aimed at completing the phylogenetic trees based on biological relevance, so OTUs representatives of at least 2 other sequences or with at least 2 reads (in total 36629 OTUs) were used in the second step and all remaining OTUs (in total 75975 OTUs) in the third step. Datasets containing the sequences were aligned and trimmed as described earlier (for both the forward and reverse datasets). Final aligned datasets had 6705 and 6691 positions in step 2 and 5845 and 5794 positions in step 3 respectively for the forward and reverse alignment. Pruned trees from the first step were used as a constraint topological tree (without branch-lengths) to infer phylogenetic trees in the second step in IQ-Tree v2.0.3 (<u>Minh et al. 2020</u>) under the GTR + Gamma + Invariant sites evolutionary model and including the ‘-fast’ option. Each phylogenetic analysis was run with 2 replicates to allow phylogenetic uncertainty, resulting in 16 total phylogenetic trees at step 2. Resulting trees from step 2 were pruned from long-branches (following step 1) and from intruders with the in-house script <u>treeCheckIntruders.py</u>’. Briefly, intruders were defined as those taxa identified under a specific supergroup and phylogenetically resolved within a monophyletic clade of a different supergroup. Final cleaned (or pruned) trees from step 2 were used as topological constraints to infer phylogenetic trees in step 3, and using the same approach for the phylogenetic inference and processing of the trees. Step 3 resulted in a total of 32 phylogenetic trees.

Total number of initial, pruned and final tips, as well as evolutionary model, constrained trees and likelihoods of every tree at every step generated and analyzed in this study can be found in **<u>Table S2</u>**. Each final tree was rooted in Discoba and Amorphea (gathering Amoebozoa, Apusomonadida, Breviatea, Holozoa and Nucletmycea) by creating an outgroup in the given node using the function ‘set_outgroup’ from the package ete3 (<u>Huerta-Cepas et al. 2016</u>); implemented in the in-house script <u>treeRootOutgroup.py</u>’).

### 6.4. Phylogenetic dating

Resulting phylogenetic trees were time-calibrated in TreePL (<u>Smith and O’Meara 2012</u>) using a total of 77 well-supported fossil calibrations across the tree (**<u>Table S1</u>**; **<u>Fig. S3</u>**). Such calibrations were chosen based on (i) monophyletic nodes, (ii) clades consistent among the 32 phylogenetic trees, (iii) at least 4 OTU representative sequences in all phylogenetic trees, (iv) consistency between branch-lengths and calibration age, and (v) the calibration has been previously validated in clade-specific molecular clock analyses. In order to allow uncertainty in calibration of the phylogenetic distance, TreePL was ran over 100 replicates on each tree, including the options ‘thorough’ and ‘log_pen’. Final dated trees were obtained with treeAnnotator v1.10.4 (<u>Drummond and Rambaut 2007</u>) by summarizing the node height to the median value. Branch lengths of the time-calibrated trees were transformed to millions of years (Ma) with the R package ‘ape’ (<u>Paradis and Schliep</u> <u>2019</u>) for following analyses. Here we have applied stringent and conservative criteria for fossil selection and calibration in order to minimize potential sources of uncertainty for each individual fossil calibration (i.e., interpretation of the fossil record and further attribution to specific nodes in the phylogenetic tree; <u>Sauquet</u> <u>2013</u>; <u>Marshall 2019</u>), yet many more fossil calibrations could potentially narrow down the posterior density.

We additionally calibrated 10 phylogenetic trees randomly chosen in a further attempt to accommodate recent findings and reinterpretations of the fossil record. The first additional calibration set (referred to as Molecular Clock 02: MC02) was chosen to include the potential early fossils *Rafatazmia* attributed to Rhodophytes at 1600 Ma old (<u>Bengtson et al. 2017</u>), used as a minimum constraint. Yet its taxonomic assignment (<u>Carlisle et al.</u> <u>2021</u>) and age (<u>Gibson et al. 2018</u>) have been questioned and therefore not considered in our primary calibration set (referred to as MC01). The second additional calibration set (referred to as MC03) was applied after the first Diatom fossils have been questioned (<u>Bryłka et al. 2023</u>), and therefore delaying the first appearance of this clade from ∼190 Ma (as a minimum constraint) to ∼125 Ma (as a minimum appearance time in the fossil record). Therefore in order to allow ambiguity in the calibration of Diatoms, the third calibration set (referred to as MC03) eliminated the crown Diatom calibration. With these additional calibrations we hope to provide a basis and a comprehensive molecular dating framework for future analyses, and further contribute to the understanding of evenly distributed calibration points over both the time and the diversity scales.

### 6.5. Lineages Through Time

As an approach to summarizing the time-calibrated trees, we computed Lineages Through Time (LTT) plots directly from the time-calibrated trees obtained in the previous section (“Phylogenetic dating”) with the in-house script ‘<u>treeLTTsubTrees.R</u>’. Given that LTT are subject to the “pull of the present” effect (<u>Nee et al.</u> <u>1994</u>) caused by the fact that extant lineages are often also young and have not had time yet to go extinct, LTT were used and only used for the purpose of summarizing complex time-trees. Supergroups were extracted with the function ‘tree_subset’ from the R package ‘treeio’ (<u>Wang et al. 2020</u>). These analyses, as well as the ones that follow (“Diversification analyses”) were carried out directly and independently on each time-calibrated tree of eukaryotic supergroups with at least 300 tips.

### 6.6. Diversification analyses

#### 6.6.1. Estimation of the diversity

Fitting diversification models to time-calibrated phylogenetic trees requires estimating the fraction of the total diversity that the analyzed tree represents. Such an estimate (hereon referred to as ‘diversity fraction’) represents the probability that a given taxa is included in the tree. However, the estimation of the total number of eukaryotic species varies dramatically among studies, ranging from 1.8-2 million species (<u>Costello et al. 2012</u>) to 5±3 million species (<u>Costello et al. 2013</u>), 8.7 ± 1.3 million species (<u>Mora et al. 2011</u>) or even up to 10¹¹ - 10¹² total microbial species (<u>Locey and Lennon 2016</u>). Far from settling this debate, here we refer to species as 99% OTUs of the 18S rDNA gene sequences avoiding any further ambiguity in the species criterium (see e.g., <u>De</u> <u>Queiroz 2007</u> or <u>Zachos 2016</u> for further details on the so-called species problem). While different species criteria cannot be directly compared, recent efforts to unify global metabarcoding data resulted in 335 662 OTUs of the V4 hypervariable region of the 18S gene (<u>Berney et al. 2023</u>) or ∼40 300 OTUs of the V9 hypervariable region of the 18S gene (<u>Singer et al. 2021</u>), agreeing with our study in a similar order of magnitude of observed OTUs (75975 OTUs of the 18S rDNA). A global survey of eukaryotic plankton diversity of the sunlit ocean (<u>de Vargas et al. 2015</u>), estimated a total eukaryotic plankton richness of ∼150 000 OTUs from ∼110 000 sampled OTUs (covering ∼73% of the total diversity). In this context, we estimated different diversity fractions to provide a minimum and a maximum scenario allowing and acknowledging ambiguity in the total number of species.

Diversity fractions were estimated on all supergroups of eukaryotes composed of at least 300 tips. We applied both the Preston’s Lognormal Model (<u>Preston 1948</u>) and the truncated lognormal model, implemented in the R package ’vegan’ (<u>Oksanen et al. 2022</u>) to search for maximum and minimum boundaries of the diversity fraction. To account for differences in abundance between the two datasets, reads were normalized at 1000, which corresponds to the closest power of ten of the smaller maximum number of reads among the two datasets.

#### 6.6.2. Rates of diversification through time

The first approach to gain insights in the early diversification of eukaryotes applies the Data Augmentation method to estimate the branch specific rates of speciation implemented in the recently developed ClaDS (<u>Maliet et al. 2019</u>) package. We used the version of ClaDS with constant turnover (turnover rate = extinction rate / speciation rate: ’ε=μ/λ’; ClaDS2) (<u>Maliet and Morlon 2022</u>). ClaDS2 was chosen because the underlying model assumptions account for rate heterogeneities across the tree, as is expected within our trees covering a wide diversity. Given that different approaches may lead to different biological interpretations (<u>Martínez-Gómez et al. 2024</u>), no other approach was applied for inferring diversification rates at such a taxonomic scale. Mean net diversification Rates Through Time (RTT; net diversification rate = speciation rate - extinction rate: ‘r=λ-μ’) and estimated number of lineages through time (or Diversity Through Time; DTT) are estimated from the reconstructed time-calibrated trees under the ClaDS module. ClaDS analyses were performed on all time-calibrated trees and both for the minimum and maximum diversity fractions. Given the size of the time-calibrated trees, certain analyses were relatively demanding in terms of computational resources and did not finish after 30 days, 4 independent attempts and up to 180 GB of RAM memory.

Supergroups such as Nucletmycea, Holozoa and Alveolata were the most affected. Completed RTT and DTT from each supergroup of eukaryotes were independently summarized into single plots by estimating the median (and 90% interquartile range) of different plots per supergroup. A summary of the resulting model parameters can be found in **<u>Fig. S12</u>**.

#### 6.6.3. Shifts in diversification rates

The second approach is aimed at identifying shifts in diversification dynamics by estimating speciation and extinction rates with the BAMM (<u>Rabosky 2014</u>) package on the time-calibrated trees, over both the minimum and maximum diversity fractions, and following the recommendations described in the online documentation (bamm-project.org/). Briefly, speciation and extinction rates are modeled within rate regimes using an exponential change function. Priors were estimated with the function ‘setBAMMpriors’ from the R package BAMMtools (<u>Rabosky et al. 2014</u>), and the control file was generated with the function ‘generateControlFile’ setting 4 MCMC chains for 10 million generations and sampled every 100 steps (for Alveolata, Holozoa and Nucletmycea MCMC chains were run for 100 million generations sampled every 1000 steps). The most plausible number of speciation rate shifts was estimated over 4 eukaryotic time-calibrated trees chosen at random. To do so, preliminary analyses were run over their extracted supergroup trees with 10 and 50 expected shifts in diversification and the posterior shift probabilities were examined with the functions ‘computeBayesFactors’ and ‘plotPrior’. For each supergroup, the most plausible number of expected shifts in configuration was chosen as the average between the highest posterior density and the highest bayes factor (**<u>Fig. S16</u>**). Final control files were generated as described above, including the corresponding estimated number of shifts. Final results were analyzed to extract the best shift configuration with the function ‘getEventData’ and ‘getBestShiftConfiguration’. Effective Sample Size (ESS) of the number of shifts was checked with the R package coda (<u>Plummer et al. 2006</u>). From the supergroups analyzed, Amoebozoa, Cryptista, Discoba, Haptista and Metamonada largely exceeded a value of 200 ESS, while the rest of the clades was above 100 ESS as a minimum. Increasing the sampling chain and/or sampling frequency would imply excessive RAM memory usage in downstream analyses (which is up to 350 GB of RAM memory for ∼4 days in some Holozoa trees). Selected shifts in speciation were summarized along the temporal dimension for each supergroup and estimated into a shift towards lower speciation rate or a shift towards a higher speciation rate.

## 7. Acknowledgements

This project was supported by the Science for Life Laboratory (SciLifeLab). MMS was partially supported by a postdoctoral fellowship from the Beatriu de Pinós programme of the Government of Catalonia’s Secretariat for Universities and Research of the Generalitat de Catalunya Economy and Knowledge (grant number: 2021BP00068) and the European Union’s Horizon Europe research and innovation programme (grant no. 101103530; MSCA postdoctoral fellowship IMAGEN3D). Computational analyses were enabled by resources in project (UPPMAX snic2020-15-340) provided by Uppsala University at UPPMAX, at the Roscoff Bioinformatics platform ABiMS (http://abims.sb-roscoff.fr; part of the Institut Français de Bioinformatique: ANR-11-INBS-0013), and the supercomputing infrastructure from the Galician Supercomputing Center (CESGA; funded by the Spanish Ministry of Science and Innovation, the Galician Government and the European Regional Development Fund; ERDF). Cedric Berney and Pierre Barbera for phylogenetic advice. Sebastian Höhna and Joëlle Barido-Sottani for comments and advice on additional diversification analysis.

## 8. Supplementary Material

**Fig. S1**: A summary of the initial constraint tree used in this study. In orange are highlighted the references used at specific nodes and in green a summary of the most relevant fossil calibrations.

**Fig. S2**: A representation at the supergroup level of the different phylogenetic trees inferred and analyzed in this study rooted at Discoba.

**Fig. S3**: All calibrated nodes over the time-tree shown in **Fig. 1A** emphasizing Proterozoic calibrations.

**Fig. S4**: Summary of median and 95% highest posterior densities of the first diversification of eukaryotes in all time-trees generated and analyzed in this study.

**Fig. S5**: Summary of the Lineages Through Time (LTT) plot of 10 randomly selected time-trees calibrated under the second calibration set (MC02). Lines represent the median among all LTT and shaded areas represent the 90% centered percentile. The gray line represents the LTT of all eukaryotes obtained from the primary calibration set (MC01) for direct comparison.

**Fig. S6**: Summary of the Lineages Through Time (LTT) plot of the different calibration sets for all eukaryotes in different tones of gray and Stramenopiles in purple.

**Fig. S7**: Diversity fractions (given in percentage) of all eukaryotic supergroups with at least 300 tips estimated by two different approaches.

**Fig. S8**: The time-calibrated phylogenetic tree shown in **Fig. 1** discriminating morphologically described OTUs (orange branches; associated to a binomial latin name, either to Genus or to Species) and environmental OTUs (blue branches; lacking morphological description). See **Fig. 1** for further details on the figure.

**Fig. S9**: Summary of the estimated Diversity Through Time (DTT) plots for the Discoba rooted scenario (upper; assuming the lowest range of diversity fractions) and the Amorphea rooted scenario (middle and lower; assuming the highest and lowest range of diversity fractions respectively). Lines represent the median, and shaded areas the median of the 90% HPD, respectively, of all independent estimated diversity through time. Gray horizontal lines represent the 90% HPD of the time scale.

**Fig. S10**: Lineages Through Time (LTT) and estimated Diversity Through Time (DTT) plots shown in **Fig. 1** and **Fig. 2**, respectively, with a focus on Archaeplastida (for the Discoba -left column- and Amporphea -right column-rooted scenarios), showing the variation through time of Chloroplastida (green algae) and Rhodophyta (red algae) and that of other main supergroups of eukaryotes for comparison. Shaded area represents the 90% centered percentile of time for the LTT plots and the 90% centered percentile of the estimated diversity for the DTT plots.

**Fig. S11**: Speciation, extinction and net diversification rates for the different rooting and diversity fraction scenarios obtained from a constant turnover rate under the cladogenetic diversification rate shift model (ClaDS, with data augmentation). Lines represent the median, and shaded areas the 90% centered percentile of the median values, respectively.

**Fig. S12**: A summary of ClaDS model parameters within all different scenarios and supergroups. Briefly, **sigma** (σ) represents the stochastic parameter of rate inheritance, **alpha** (α) the deterministic trend in speciation rates changes, the **trend component** (*m*=α×exp(σ^2^/2)) the average trend at speciation (whether daughter rates tend to be higher or lower than parental rates), **epsilon** (ε) the extinction rates at constant turn-over, **Lambda0** (λ0) the speciation rate at the beginning of the process, and **Lambda-tip** (λtip) the median speciation rate at present-day (tips).

**Fig. S13**: Total number of speciation rate shifts up (solid colours) or down (shaded colours) for all supergroups by geological period across the different 32 time-trees. The cartoon on the top left box exemplifies how the shift count has been obtained.

**Fig. S14**: Proportion of shifts in the speciation rate (square rooted) across the different geological periods, for the Discoba rooted scenario and the Amorphea rooted scenario, accounting for the maximum diversity fraction and the minimum diversity fraction. ‘Growth’ indicates a shift towards a higher speciation rate and ‘decay’ indicates a shift towards a lower speciation rate. The proportion of shifts was calculated by dividing the total number of shifts per time period (**<u>Fig. S13</u>**) by the total branch-length within the given time period.

**Fig. S15**: A graphical representation on the estimation of the diversification of the Last Eukaryotic Common Ancestor (LECA) from different studies cited herein. Blue vertical rectangles correspond to the range estimated by our analyses, given by the black bars (the dot represents the median).

**Fig. S16**: Bayes factor and most frequent number of the observed number of shifts in speciation rates along the MCMC run from the BAMM analyses of all supergroups analyzed in this study.

**Table S1**: List of all fossil calibrations used in this study.

**Table S2**: A summary of the number of branches in each step of the phylogenetic tree building and the likelihood of the trees.

