## Supplementary figures and images for "Environmental phylogenetics supports a steady diversification of crown eukaryotes starting from the mid Proterozoic"

### Fig. S1

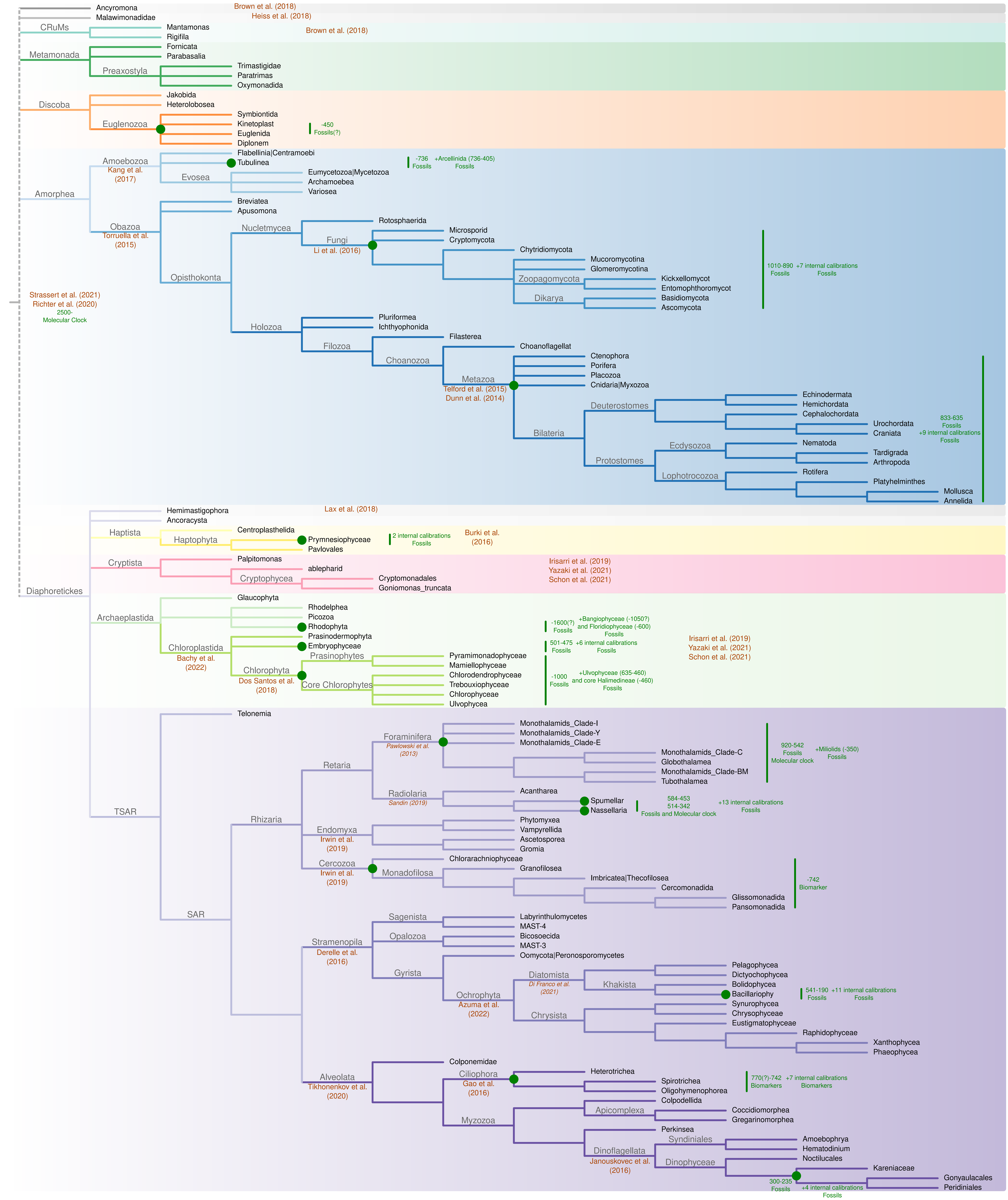

### Fig. S3

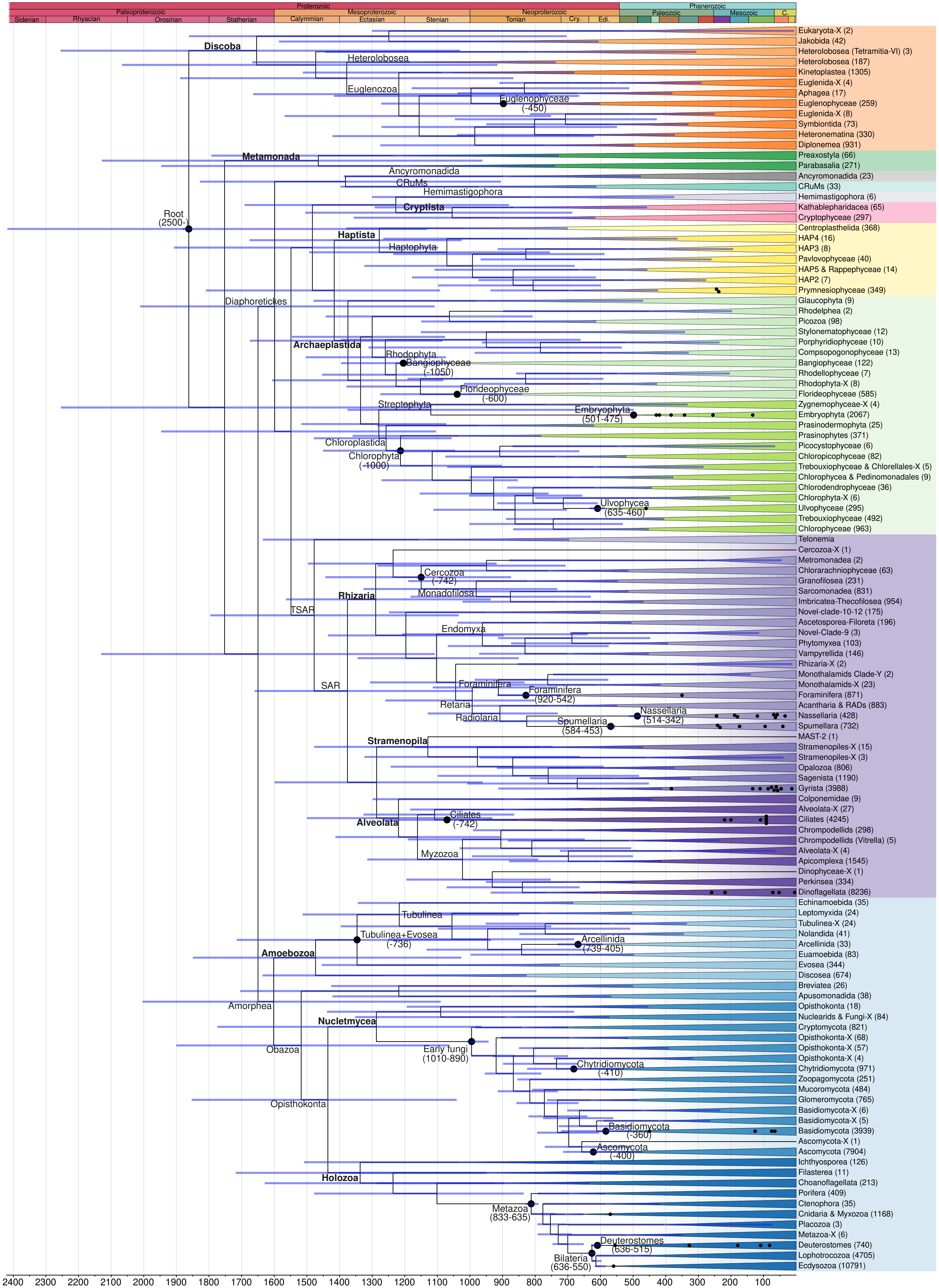

### Fig. S4

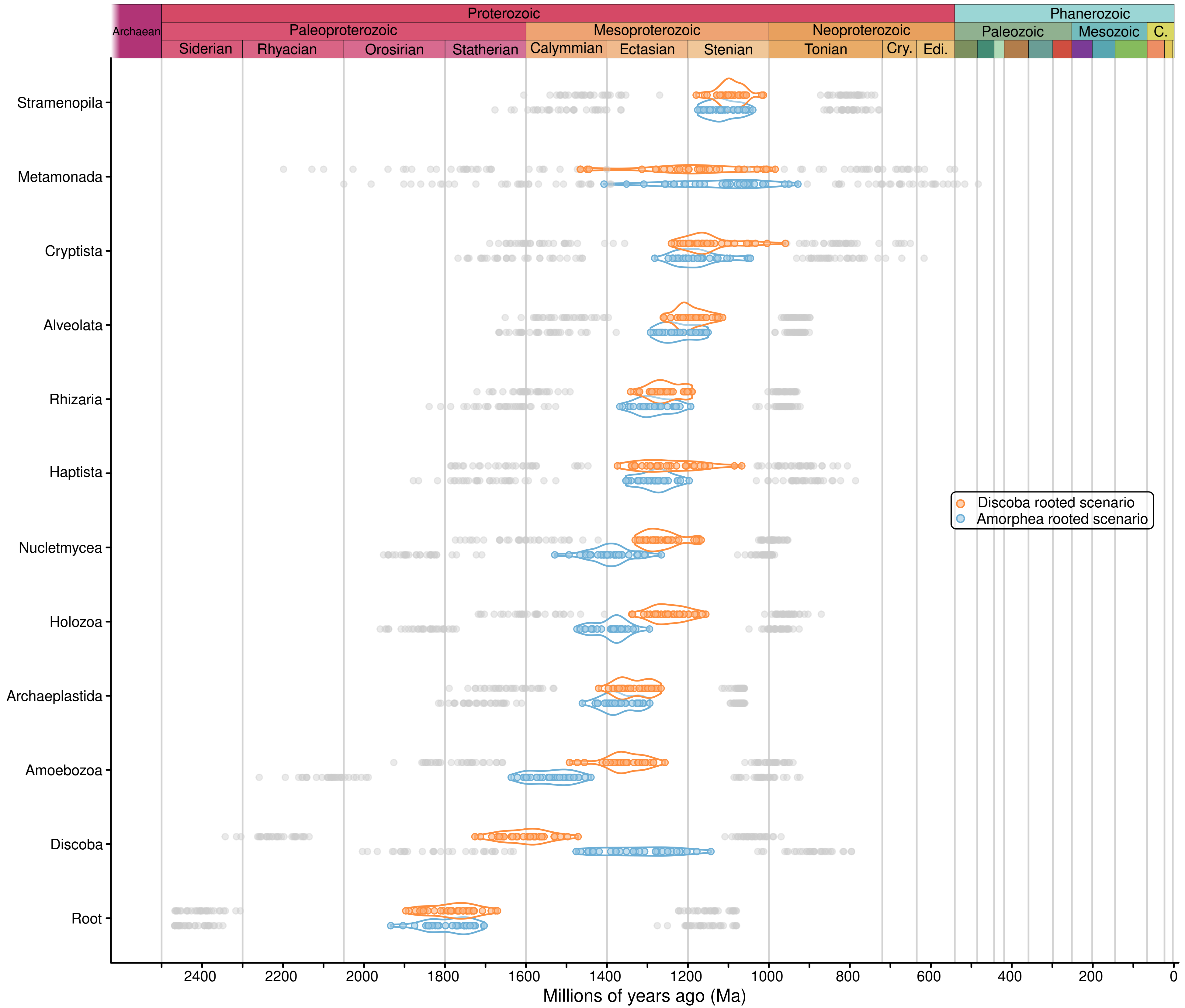

### Fig. S5

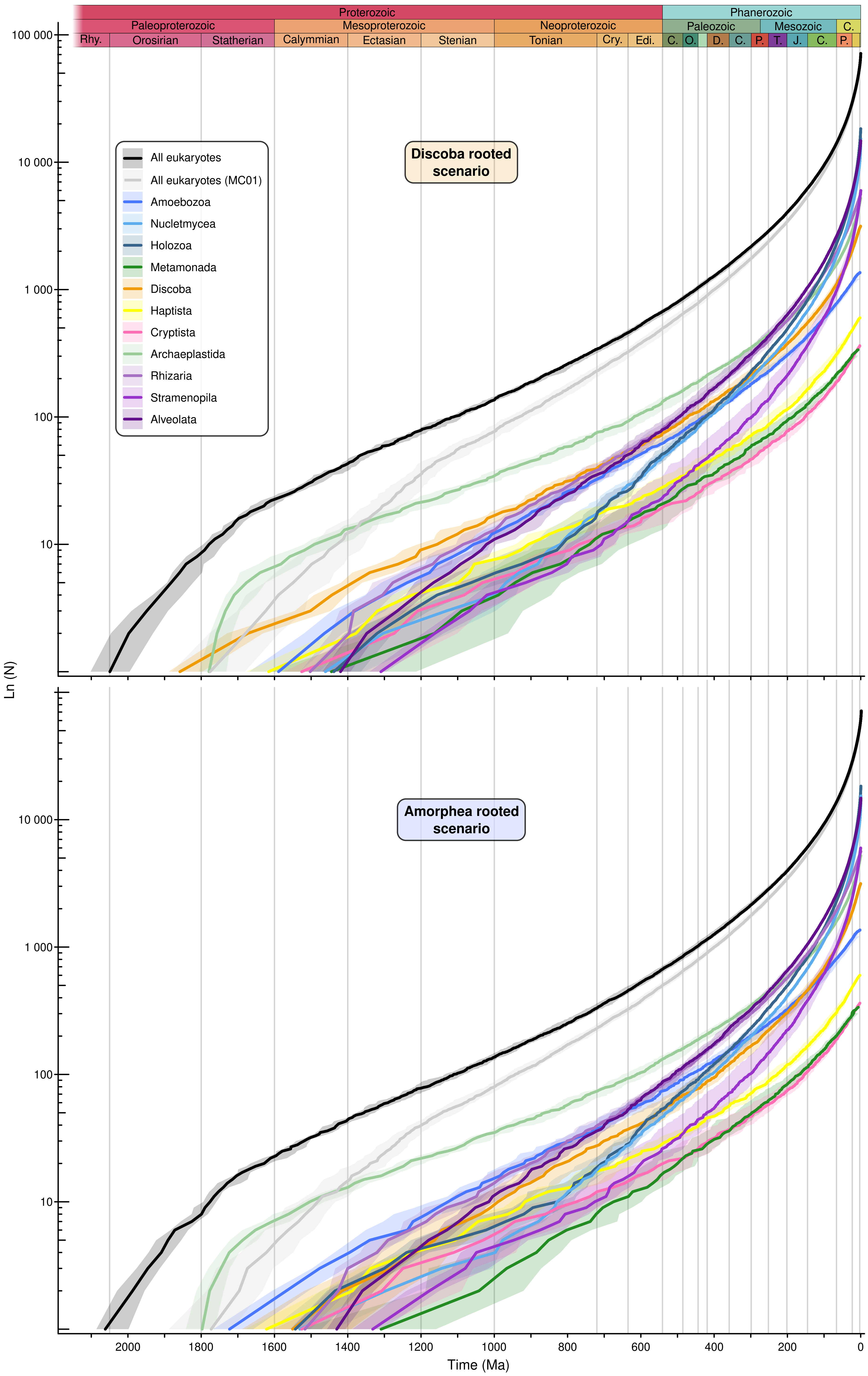

### Fig. S6

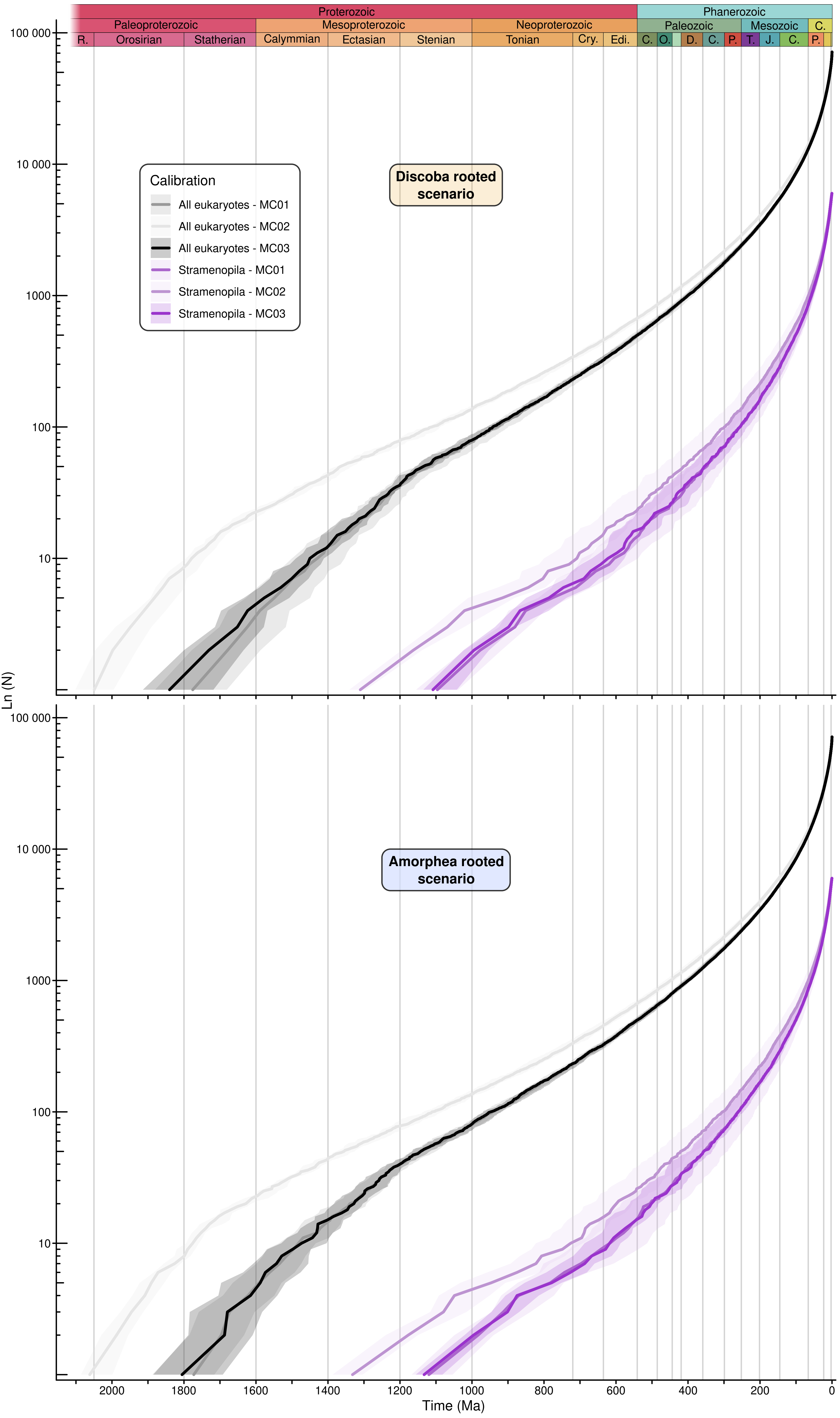

### Fig. S7

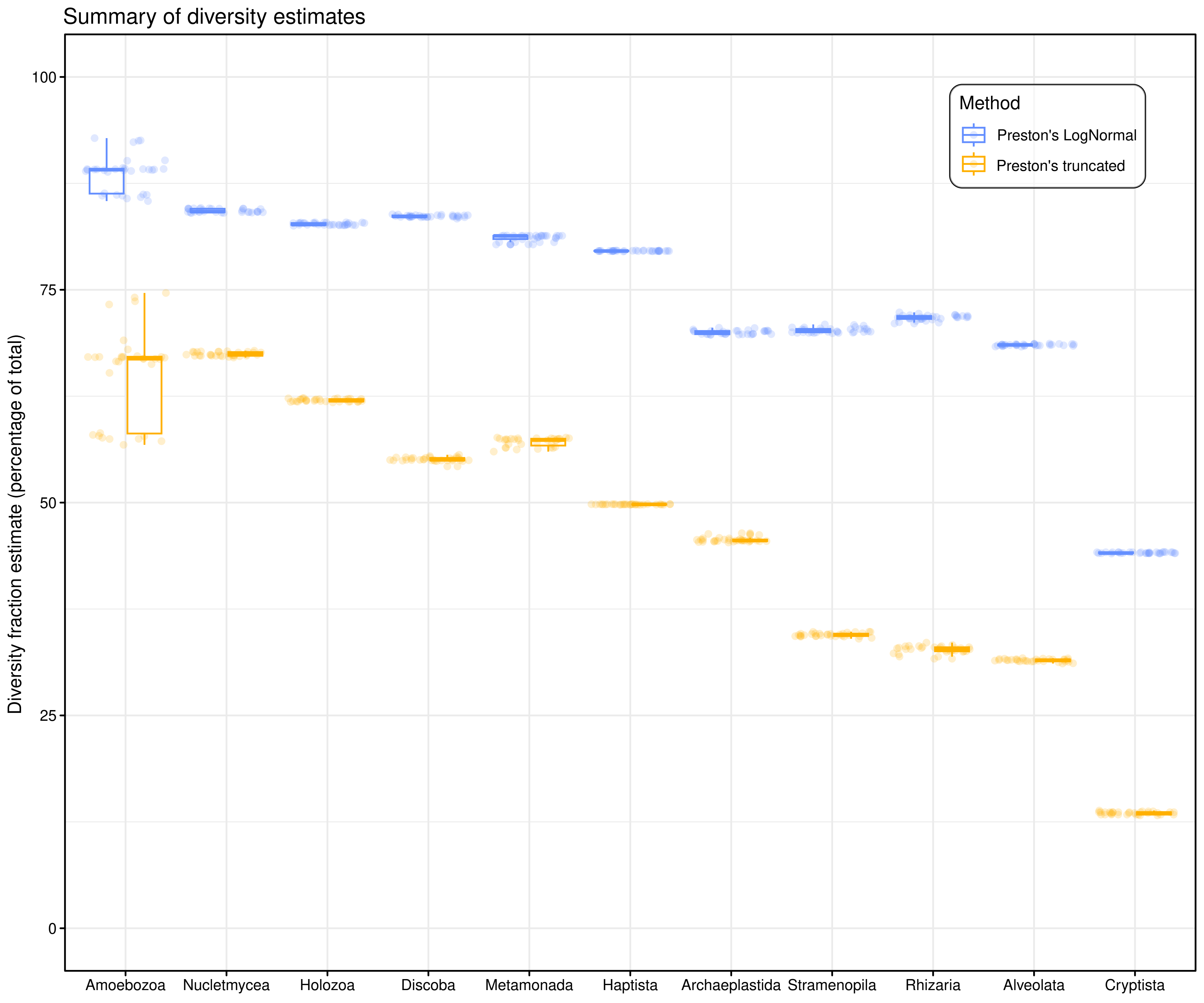

### Fig. S8

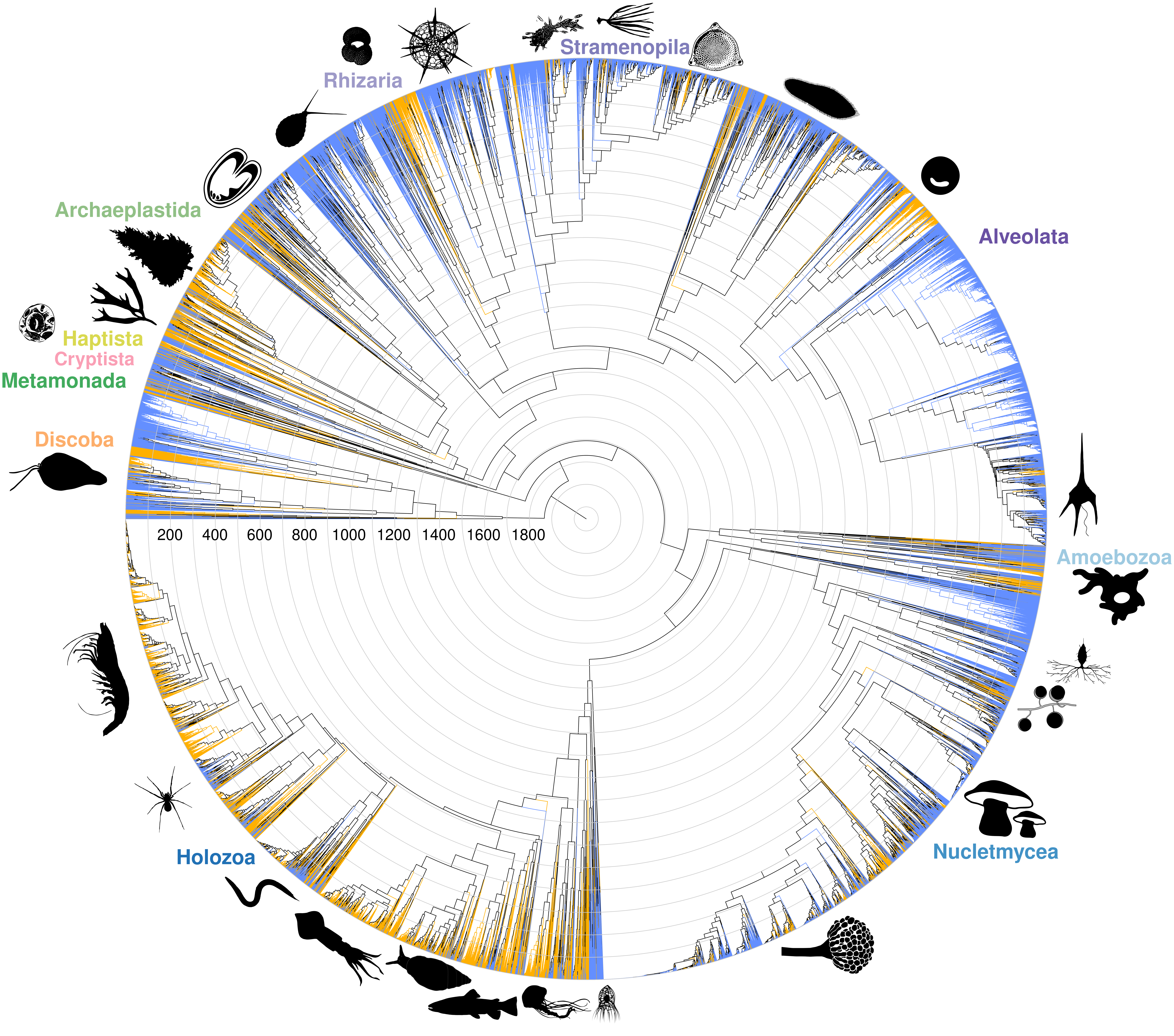

### Fig. S9

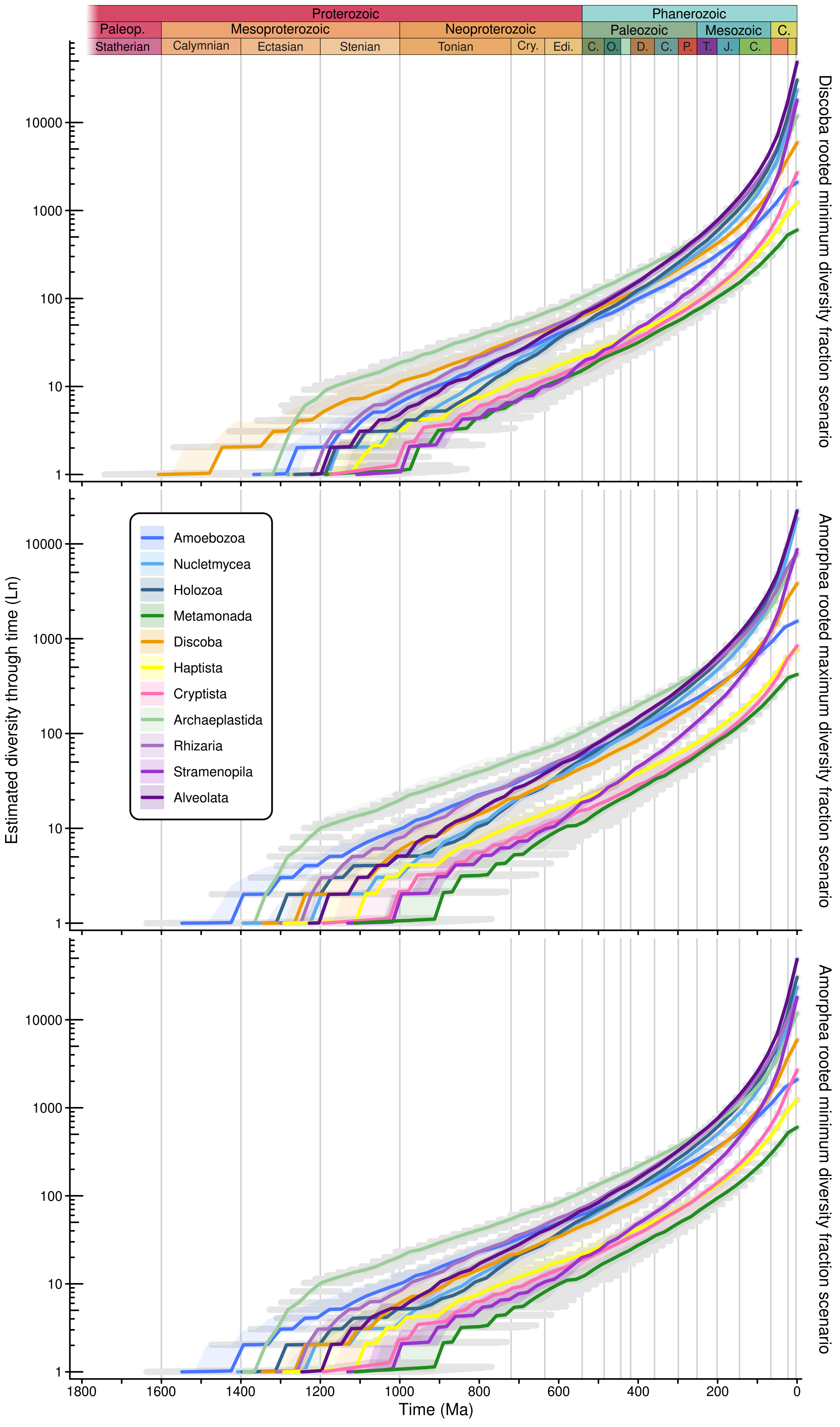

### Fig. S10

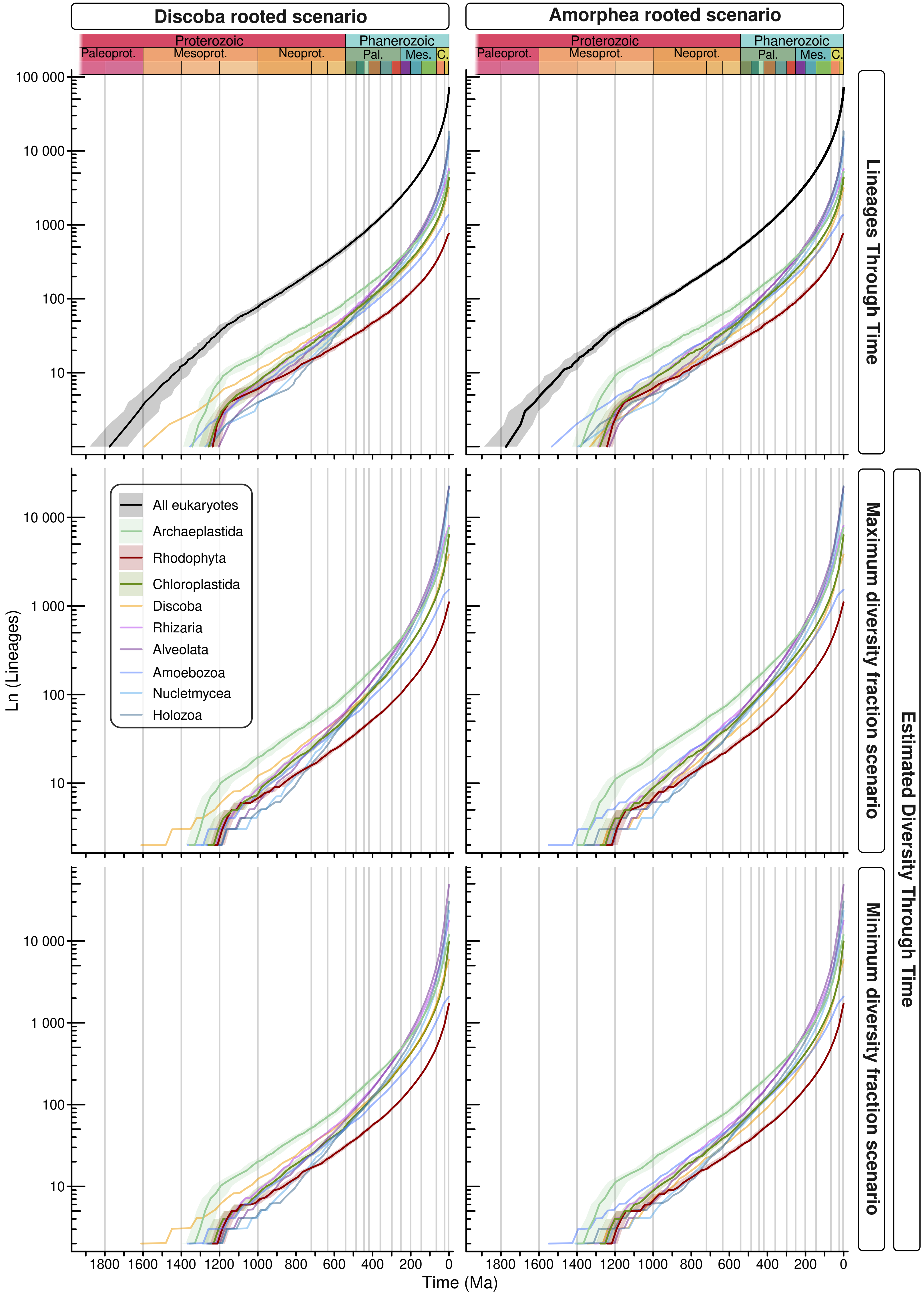

### Fig. S11

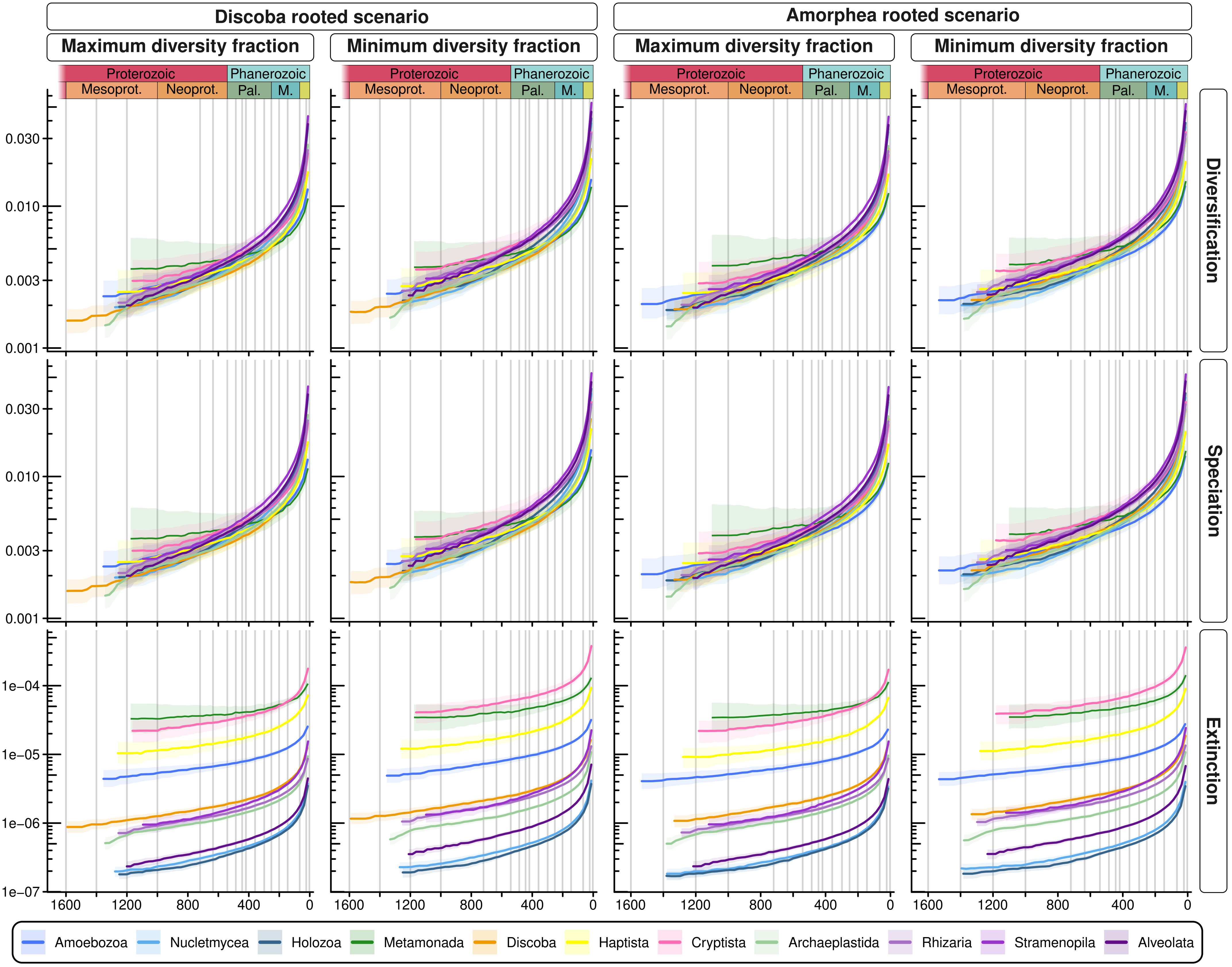

### Fig. S12

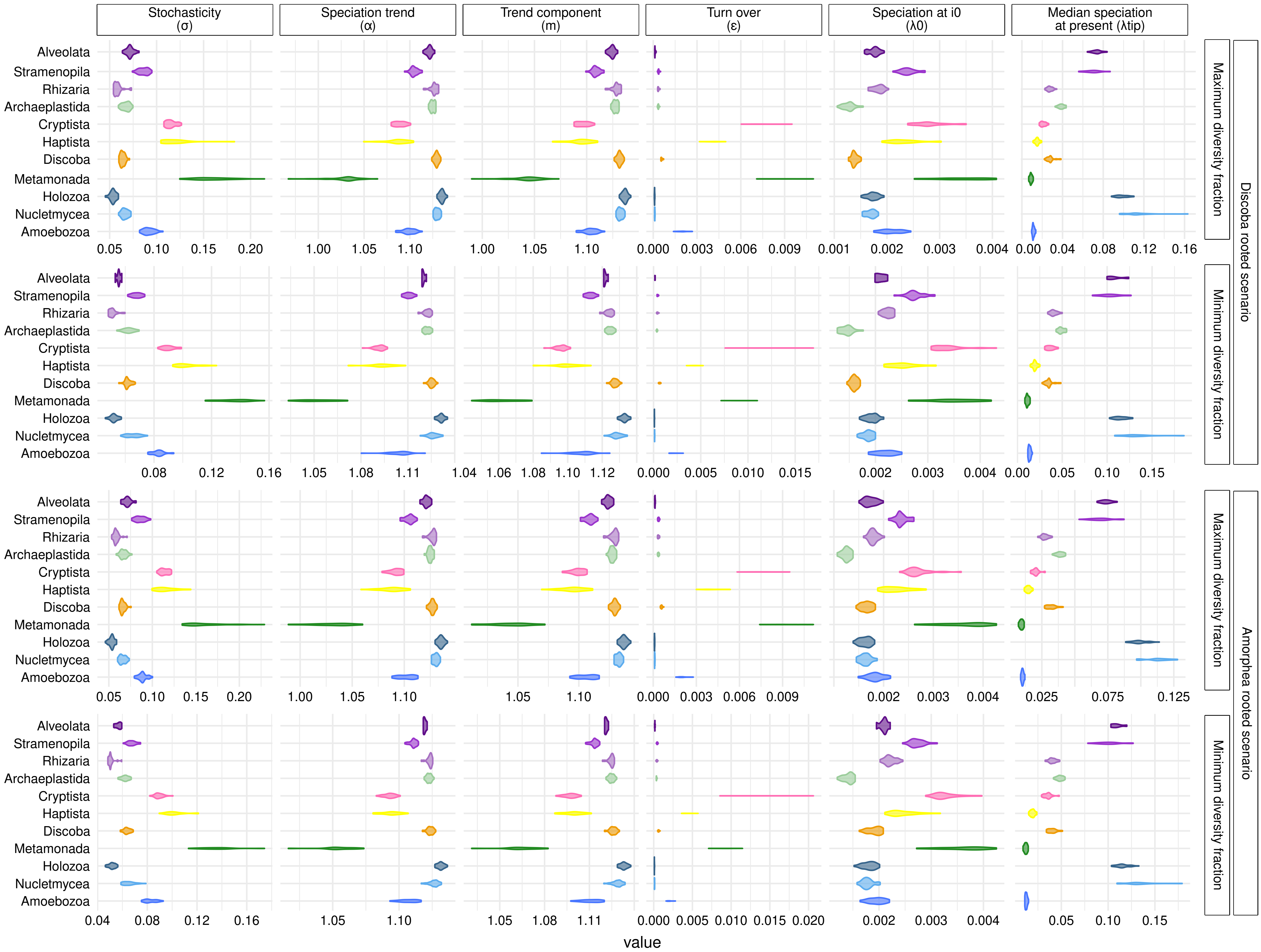

### Fig. S13

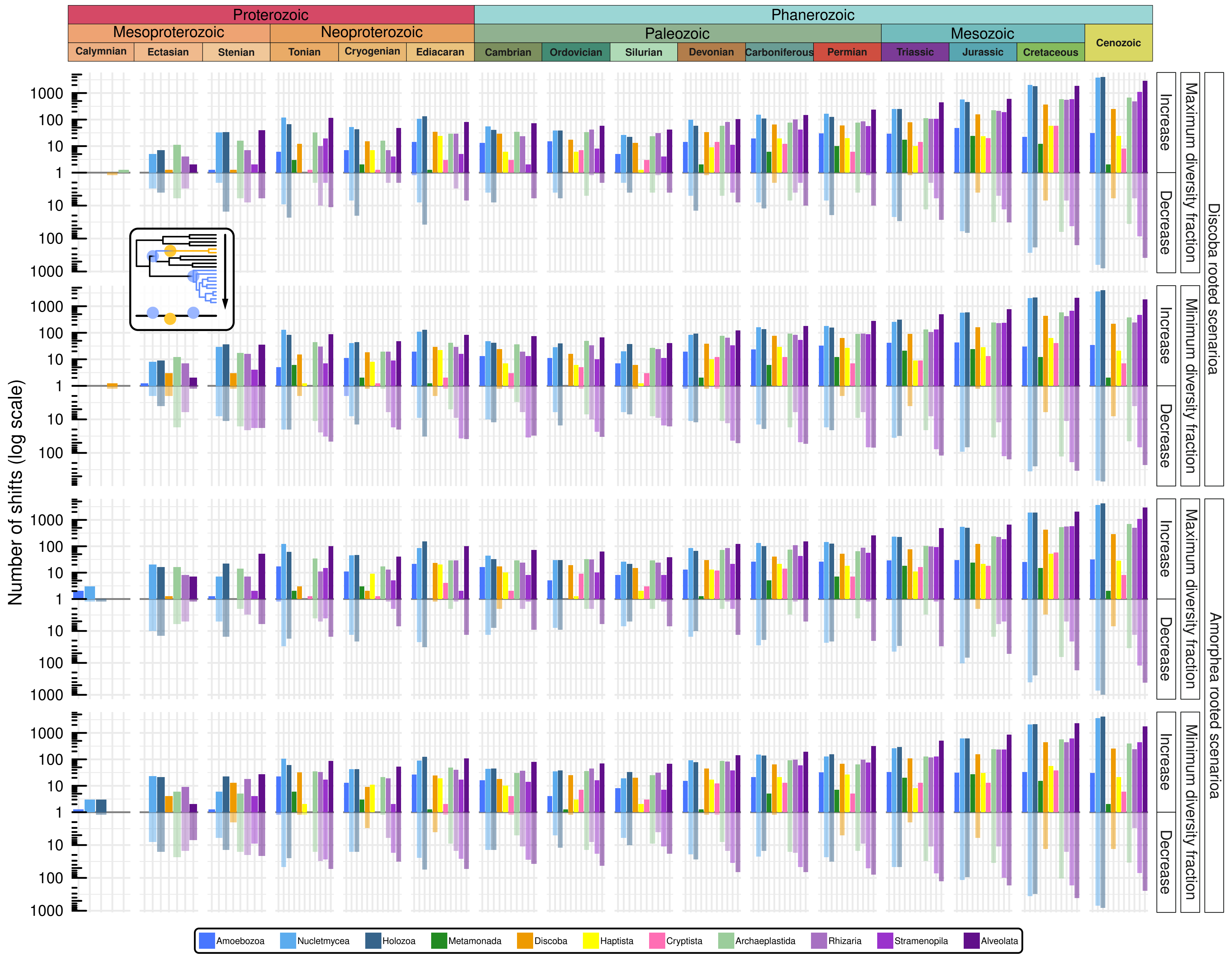

### Fig. S14

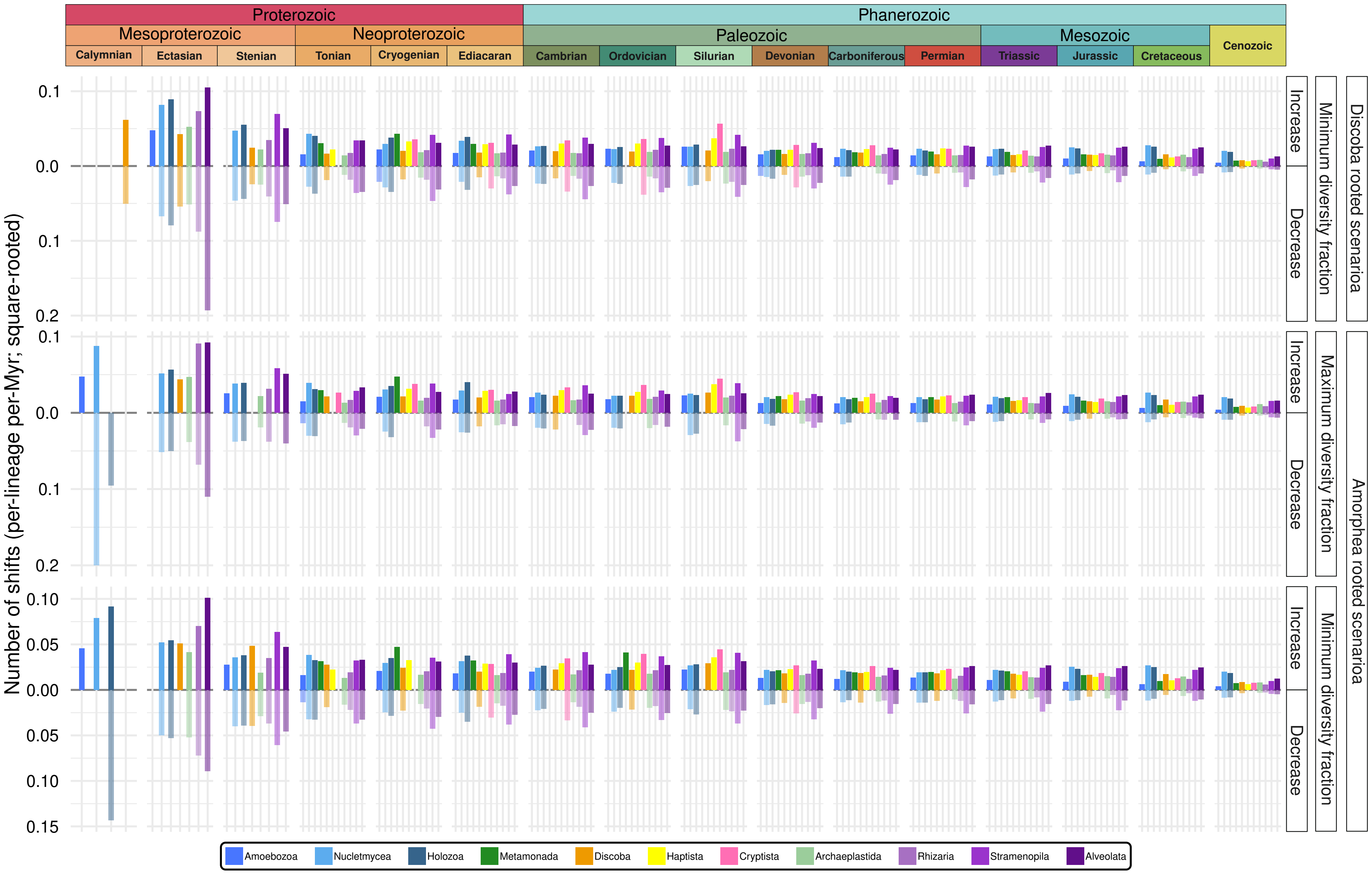

### Fig. S15

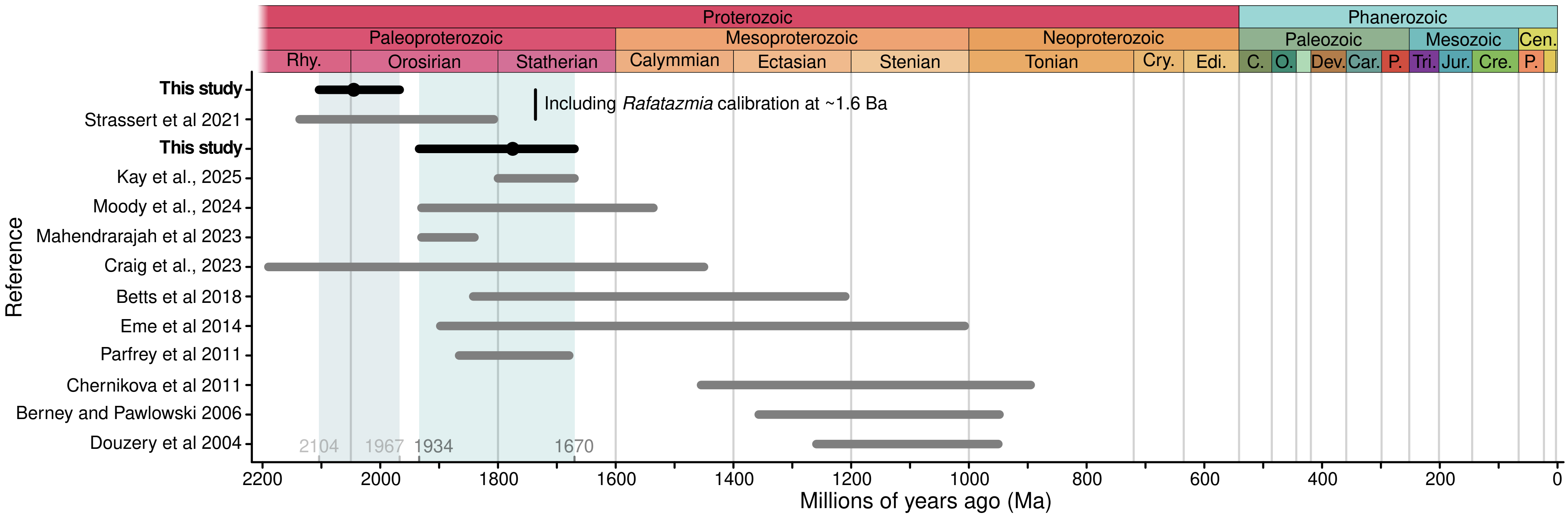

### Fig. S16

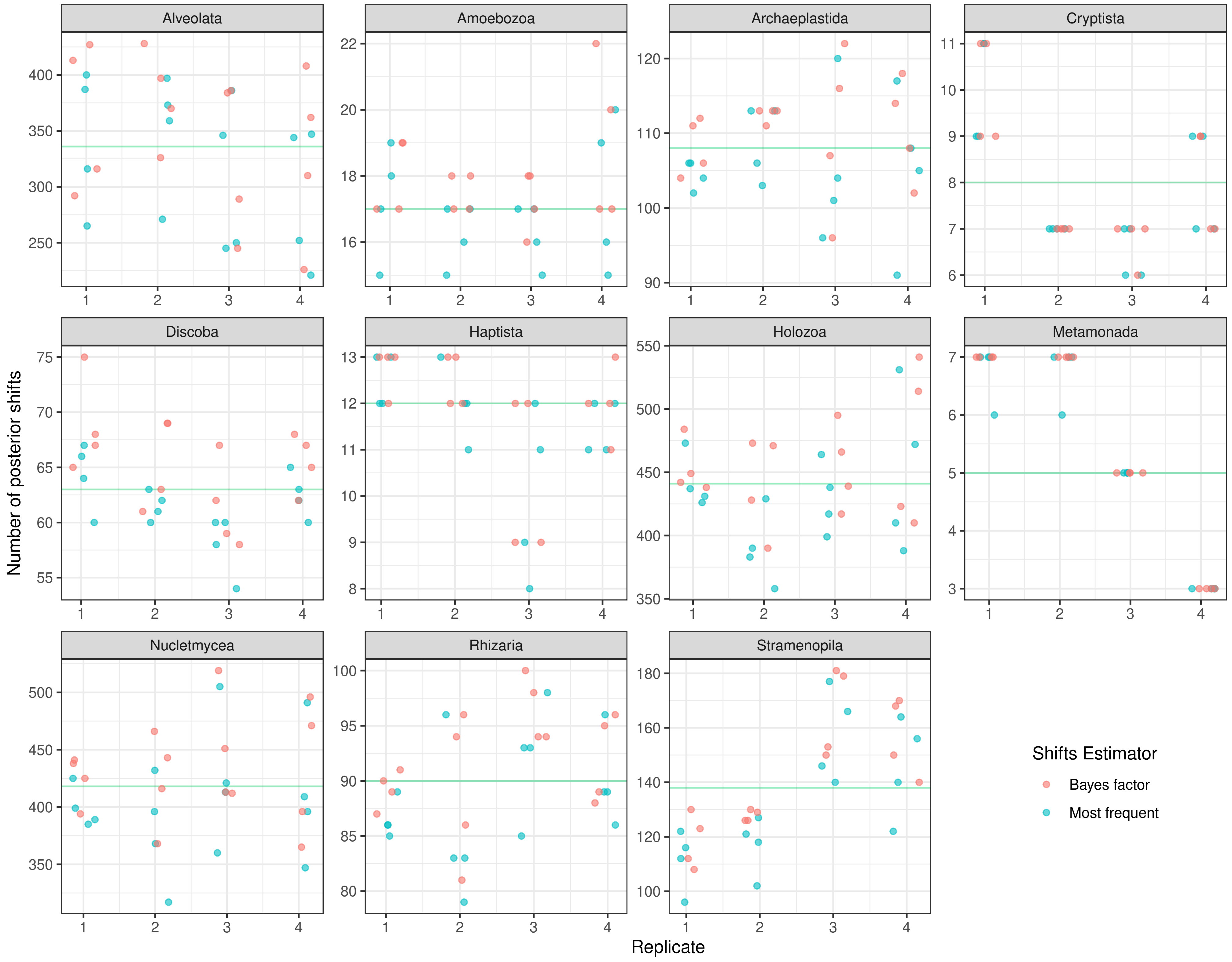
