## Supplementary material for "Environmental phylogenetics supports a steady diversification of crown eukaryotes starting from the mid Proterozoic": Fig. S2

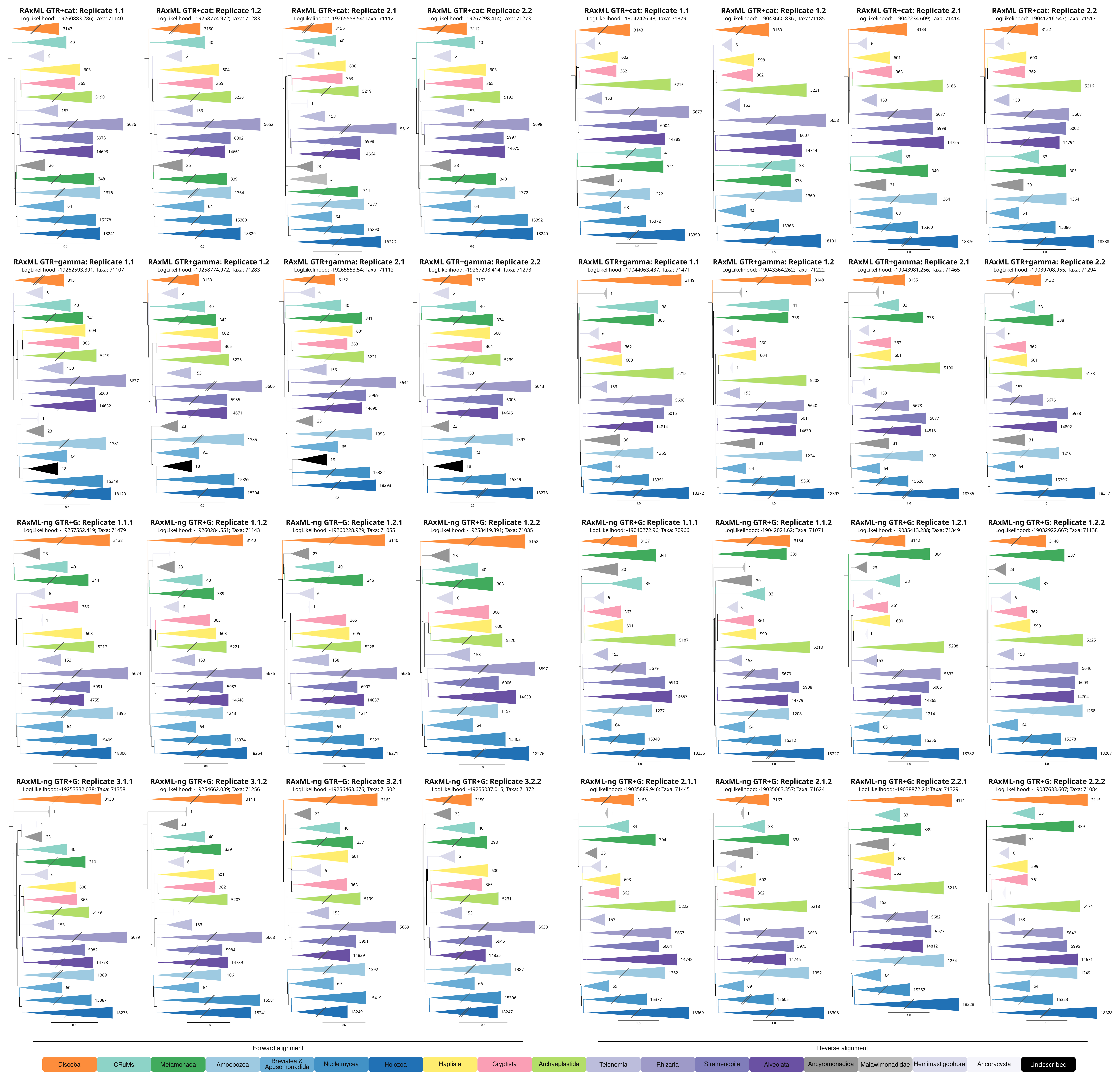


**Figure S1**. Overall topology of the different phylogenetic analyses. Titles above each tree refer to the software and model used to build the phylogenetic tree at step 1 (including OTUs with at least 10 reads), thereafter, sequences were added progressively using the previous tree as constraint (first OTUS > 1 reads, then OTUs with 1 read) using the fast option of IQtree. Resulting phylogenetic trees were pruned from (i) long branches (defined by outliers from a normal distribution) and intruders (if a sequence associated taxonomy falls within a different supergroup node). Numbers after clades (triangles) represent number of sequences of every clade. Clades with a double barred symbol are 25% reduced and single barred symbols are 50% reduced for clarity.
